# Tissue mechanics sets developmental scaling of collective cell migration

**DOI:** 10.64898/2026.08.19.745692

**Authors:** Boris Guirao, Aurélien Villedieu, Julien Delpierre, Fabian Gärtner, Lale Alpar, Isabelle Gaugué, François Graner, Floris Bosveld, Yohanns Bellaïche

**Affiliations:** Institut Curie PSL Research University, CNRS UMR3215, INSERM U934; Paris Cedex 05, F-75248, France; Sorbonne Universités, UPMC Univ Paris 06, CNRS, CNRS UMR3215, INSERM U934; Paris, F-75005, France; Université Paris Cité, CNRS, MSC UMR 7057; Paris, F-75006, France

**Author notes:** These authors contributed equally to this work (Boris Guirao, Aurélien Villedieu, Julien Delpierre). These authors contributed equally to this work (Floris Bosveld, Yohanns Bellaïche). Institut Pasteur, Université de Paris, CNRS UMR3738, Developmental and Stem Cell Biology Department; Paris, F-75015, France. co-corresponding authors;, **Correspondence and requests for materials** should be addressed to Y.B.

## Abstract

Despite substantial variation in adult size, animals within a species maintain consistent tissue patterns and shapes, a core property known as developmental scaling or size invariance. Developmental scaling has predominantly been attributed to the scaling of morphogen gradients and gene patterns to maintain positional information and cell fate specification^1,2^. However, development also necessitates collective cell flows that reshape tissues and reposition cells^3,4^. How these flows adapt to body size remains unclear. By combining quantitative live imaging, experimental perturbations, and physical modeling in the *Drosophila* thorax epithelium, we address this question in the context of a fundamental process: collective cell migration. We find that migration velocity scales linearly with tissue size, accounting for size-invariant cell positioning. While gene patterning scales with tissue size and modulates force generation, it is not sufficient to ensure proper velocity scaling. Instead, tissue mechanical properties govern the dependence of migration velocity on tissue size, enabling developmental scaling within the physiological range of animal sizes. These findings uncover principles and limits of size invariance by revealing how tissue mechanics sets the scaling behavior of collective cell flows with organismal size.

---

Animals display an extensive range of sizes, a physiological variation that is readily apparent when comparing different species but also within a given species. Despite these size differences within a species, animals maintain consistent cell allocations and body proportions essential for function and survival. Much effort has been devoted to understanding how differences in growth during development, including changes in growth rates and the duration of growth phases, contribute to variation in organism size^5–7^. Furthermore, building on seminal experiments demonstrating the scaling of embryonic development in half-sized embryos, numerous studies have delved into how the size of morphogenetic fields and, more specifically, gene patterns scale in relation to tissue or animal size^1,8^. These investigations have been pivotal in elucidating how morphogen gradients adapt to tissue size to specify the correct cell fate distribution during development^9–20^. However, tissue development also necessitates the generation of forces that drive tissue flows, thereby controlling cell positions or modulating animal shape and allometry^3,4^. The mechanisms governing the scaling of these collective cell flows have received substantially less attention, impeding our understanding of how tissue development adapts to animal size.

Collective cell migration is a major developmental process also involved in wound healing and cancer invasion^21,22^. Drawing on *in vivo* and *in vitro* systems, extensive work has investigated the mechanisms controlling migration velocity, directionality, and the basis of cell-cell cohesiveness^23–25^. While numerous molecular and mechanical signals trigger and orient collective cell migration, displacement can be propelled by supracellular actomyosin cables, cell-polarized protrusive activity, and the generation of traction forces on the extracellular matrix (ECM)^26–38^. Additionally, cell-cell or cell-ECM interactions and global tissue polarization signals play crucial roles in coordinating collective cell migration^29,30,36,39–45^. Complementarily, theoretical models of collective cell migration have explored how traction forces, cell-cell interactions and tissue mechanical properties explain collective cell dynamics^28,31,43,46–53^. Despite these advancements, we still lack an understanding of whether and how the process of collective cell migration adapts to animal size to ensure proper cell positioning and tissue organization.

Here, after first establishing the contribution of collective cell migration to epithelial morphogenetic tissue flows during *Drosophila* pupal development, we investigate how this migration is adapted to animal size. Our findings reveal that while the duration of migration remains constant, the velocity of collective migration linearly increases with tissue size, a sufficient condition to ensure robust cell positioning within the animal. Through a combination of experimental approaches and theoretical modelling, we show that while gene patterning scales with tissue size, tissue mechanical properties enable the emergence of a linear scaling of velocity within the physiological range of animal sizes. This work uncovers new limits in the scaling of tissue flows despite the scaling of morphogenetic field size. Together, our findings highlight the respective contributions of tissue patterning and mechanical properties in the adjustment of collective cell flow to animal size.

## Thorax flow velocity shows a reproducible pattern and scales with tissue length

As described in prior studies, the development of the *Drosophila* dorsal thorax epithelium (notum) can be visualized via confocal time-lapse microscopy, using the adherens junction (AJ) marker Ecad:3xGFP^54–58^ (Fig. 1a-b and Supplementary Video 1). Such observations have unveiled a collective anterior-directed flow of thorax cells during metamorphosis^55,58,59^. To begin to probe the correlation between tissue size and flow, we first characterized flows in notum tissues of commensurate lengths (between 616 and 707 µm, with mean tissue length *L*_*ref*_ = 656 µm) from 15 to 40 hours after pupa formation (hAPF) within a medial region of the tissue (Fig. 1b). Upon spatial and temporal registration of each time-lapse movie, tissue flows in each animal were quantified employing Particle Image Velocimetry (PIV, Fig. 1b and Methods). As anticipated, the tissue flow is primarily aligned with the antero-posterior (AP) axis (taken as the horizontal *X* axis, Fig. 1b). Given the homogeneity of the velocity field along the medio-lateral (ML) axis (Fig. 1b), the AP velocity was first averaged across the ML axis. Then, it was averaged across animals at each normalized AP position *x* (*x* = *X*/*L*, where *L* is the tissue length, Fig. 1b), thereby enabling us to plot the averaged velocity profile along the tissue AP axis as a function of developmental time (Fig. 1c). This kymograph of the flow velocity effectively captures the AP dynamics of thorax flow in space and time, highlighting its AP heterogeneity with higher velocities in the anterior region of the notum. Tissue flows start around 19 hAPF and end at 36 hAPF, exhibiting an acceleration phase from 19 to 25 hAPF, and a deceleration phase from 28 to 36 hAPF, reaching peak AP velocities around 25 hAPF (Fig. 1c, Extended Data Fig. 1a and Supplementary Video 1). In what follows, we thus considered the mean velocity between 24 and 28 hAPF (*V*), which reaches a maximum *V*_*max*_ = 9.9 ± 1.0 µm/h at *x* = 0.9 (Fig. 1d).

**Fig. 1.**
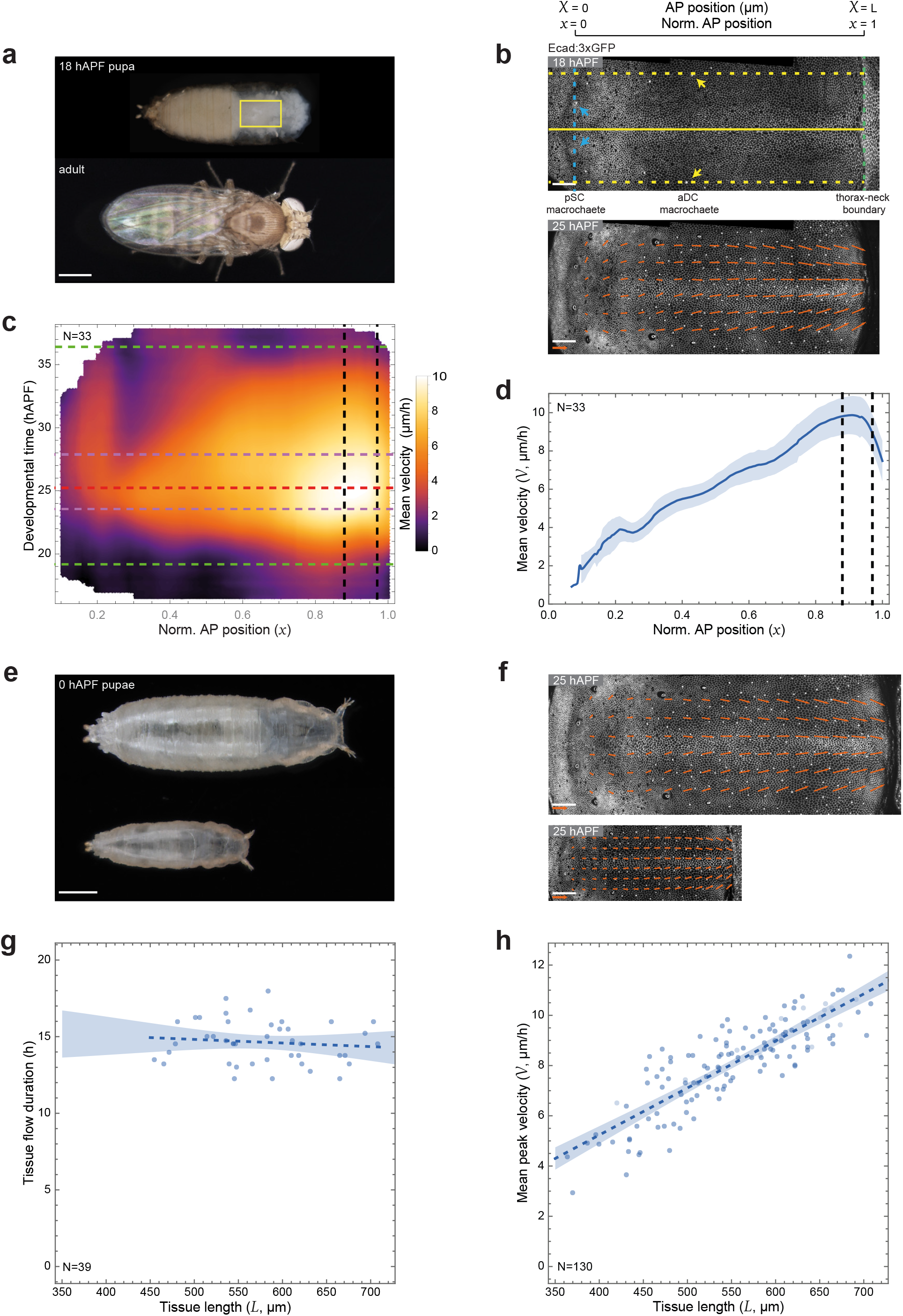
Tissue flow velocity scales with tissue length. The AP axis is horizontal with the posterior to the left. Unless otherwise stated: solid line, mean experimental data; dark blue dashed line, linear fit; shaded region, SD. N, number of animals. Genotypes are listed in Extended Data Table 3. *x* = *X*⁄*L*. **(a)** Images of a *Drosophila* pupa at 18 hAPF with the pupal case removed from head and thorax (top) and an adult (bottom). Yellow rectangle, ROI for tissue flow analyses (b). **(b)** Ecad:3xGFP notum (*L* = 673 µm) live images at 18 (top) and 25 (bottom) hAPF. In the top image arrows indicate landmark macrochaeta (yellow, anterior dorsocentral, aDC; blue, posterior scutellar, pSC). The ROI for PIV analyses is defined by the intersecting blue dashed line (pSC position along AP), yellow dashed lines (aDC position along ML) and the green dashed line (thorax-neck boundary). The blue and green dashed lines set the AP (*X* = 0, *X* = *L*) and normalized AP (*x* = 0, *x* = 1) positions, respectively. Yellow solid line, midline. Arrows in the bottom image show the local flow velocity averaged from 24 to 28 hAPF. **(c)** Kymograph of the mean velocity (averaged across tissues) along the normalized AP position (*x*) as a function of developmental time, for commensurate tissue lengths (616 < *L* <707 µm). Green dashed lines, tissue flow onset (19 hAPF) and end (36 hAPF) as defined by 20% of the *V*_max_ (see Extended Data Fig. 1a). Magenta dashed lines, peak of velocity (23.6 and 27.9 hAPF), as defined by 90% of the *V*_max_ (see Extended Data Fig. 1a). Red dashed line, *V*_max_ (25.2 hAPF). Vertical black dashed lines, peak of velocity along the AP axis (*x* = 0.88-0.97). **(d)** Graph of the mean velocity (*V*, averaged from 24 to 28 hAPF) as a function of the normalized AP position (*x*) for commensurate tissue lengths (616< *L* <707 µm). Vertical black dashed lines, peak of velocity along the AP axis (*x* = 0.88-0.97). Same data as c. **(e)** Images of a large and a small pupa at 0 hAPF. **(f)** Ecad:3xGFP notum live images at 25 hAPF with local flow velocity averaged from 24 to 28 hAPF (arrows) for *L* = 726 (top), and *L* = 345 µm (bottom). **(g)** Graph of the tissue flow duration as a function of tissue length (*L*). Duration for each animal was defined as the interval between the two time-points for which the tissue velocity is above 20% of *V*_*max*_ (see Extended Data Fig. 1a). Shaded area, 95% CI of the mean. R^2^= 0.012, *p*= 0.478 (Student’s *t*-test). Includes all data from c. **(h)** Graph of the mean peak velocity (*V*) (velocity averaged from 24 to 28 hAPF and over 0.88 < *x* < 0.97) as a function of tissue length (*L*). Shaded area, 95% CI of the mean. R^2^= 0.72, *p*= 2.94 × 10^−37^ (Student’s *t*-test). Includes all data from g. Scale bars: 1 mm (a, e), 50 µm (b, f), and 10 µm/h (arrows b, f).

Animal size varies significantly in *Drosophila*, primarily due to sexual dimorphism and nutrition during the larval stages prior to metamorphosis^60,61^ (Fig. 1e). To investigate the correlation between tissue flows and animal size, we analyzed animals of both sexes across a wide size range at the onset of metamorphosis. In total, we imaged 130 animals (61 females and 69 males) with notum lengths ranging from 364 to 707 µm. The systematic spatiotemporal characterization of thorax flows by PIV enabled us to assess whether the duration or velocity of tissue flows was influenced by tissue size and sex (Fig. 1f and Supplementary Video 2). Although the total duration of thorax flow did not change with the tissue length (Fig. 1g and Extended Data Fig. 1b), strikingly, the peak value of tissue flow velocity linearly increased with tissue length for both males and females (Fig. 1h and Extended Data Fig. 1c). This linear scaling can also be observed along the AP axis (Extended Data Fig. 1d-f). We therefore defined for each animal its rescaled velocity 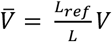, namely the AP velocity rescaled by tissue length, where *V*/*L* is multiplied by *L*_*ref*_. As expected, the kymograph of the rescaled velocity across 130 animals exhibited a smaller standard deviation at each AP position and developmental time than that of the velocity (Extended Data Fig. 1g-j). Collectively, these data establish that thorax epithelial flow velocity linearly scales with tissue length.

### Thorax flow is associated with an active migration on the apical extracellular matrix

To investigate how tissue flow velocity scales with tissue length, we first sought to understand the forces underlying the thorax flow velocity profile. Tissue flows can result from pulling or pushing forces generated by an active neighboring morphogenetic event^62,63^. The thorax flow is associated with the anterior invagination of the neck^59,64^, potentially providing a pulling force on the epithelium. To mechanically uncouple thorax flow from this morphogenetic event, we used repeated laser ablations to cauterize the neck tissue prior to invagination and prevent wound healing (Fig. 2a). We then recorded tissue flow to quantitatively assess the rescaled velocity along the AP axis. As expected, ablation immobilized the adjacent tissue, which behaved as a physical obstacle (Supplementary Video 3). Nonetheless, more posteriorly from the ablated region, thorax flow proceeded at velocities comparable to non-ablated animals, indicating that most thorax flow is independent of neck invagination (Fig. 2b and Supplementary Video 3). Next, we performed repeated ablations in the posterior notum to test whether a pushing force could drive the anterior thorax flow (Fig. 2a). Posterior ablation also led to immobilization of the surrounding tissue. Yet, thorax flow in the most anterior region exhibited dynamics similar to those of control tissue (Fig. 2b and Supplementary Video 4). Altogether, these data indicate that thorax AP flow is minimally affected by remote anterior and posterior morphogenetic events.

**Fig. 2.**
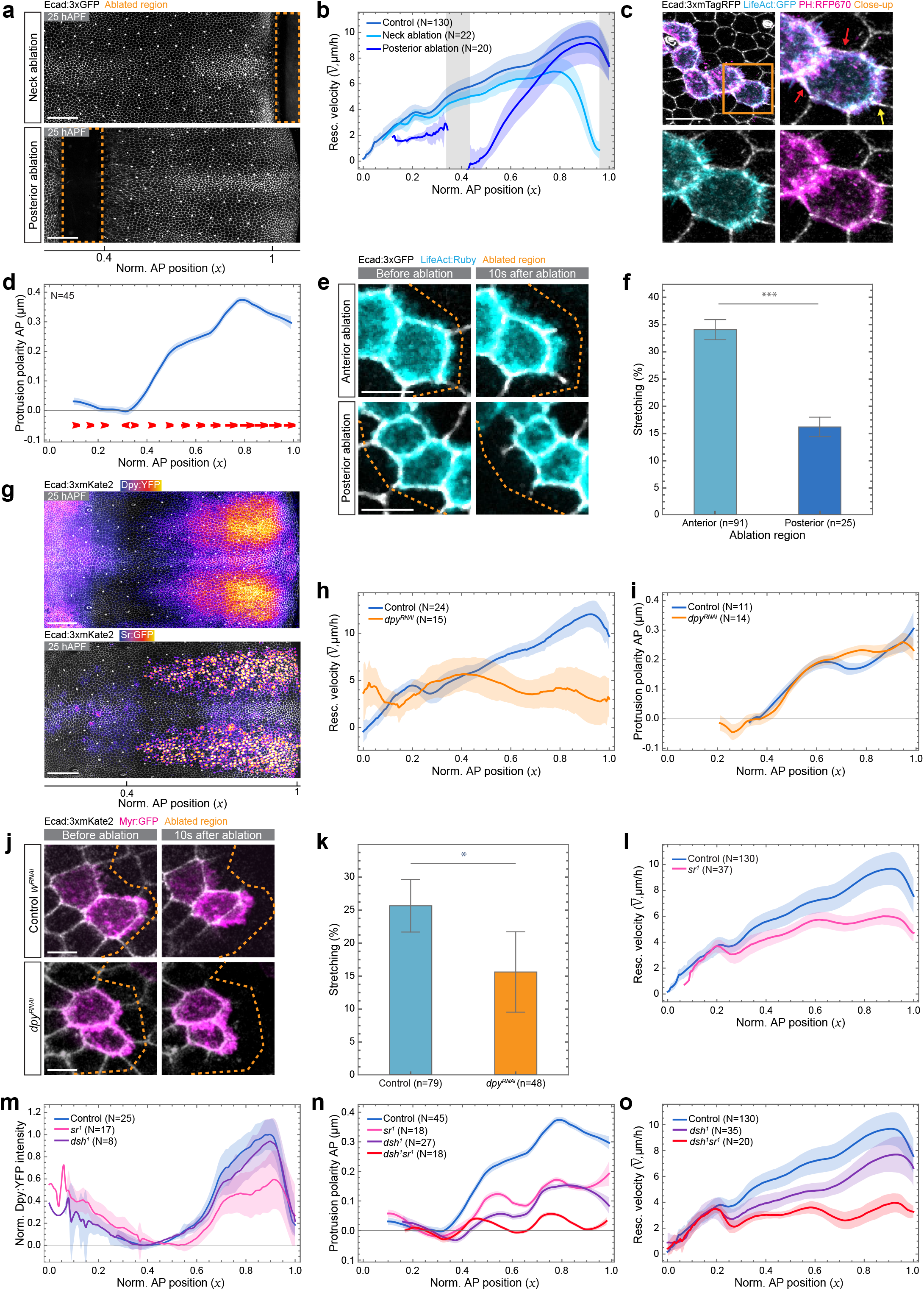
Thorax flow relies on a collective migration on the apical extracellular matrix. The AP axis is horizontal with the posterior to the left. When relevant, the normalized AP position (*x*) is indicated below the images. Images were acquired in the anterior notum region at 25 hAPF. Quantifications are averaged from 24 to 28 hAPF. Unless otherwise stated: solid line, mean experimental data; shaded region, SD. N, number of animals. n, number of ablations. Genotypes are listed in Extended Data Table 3. 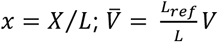. **(a)** Ecad:3xGFP notum live images after neck ablation (top, *x* = 1) and ablation posterior to the aDC macrochaetae (bottom, *x* = 0.4). Orange dashed boxes, ablated regions. **(b)** Graph of the mean rescaled velocity 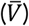 as a function of the normalized AP position (*x*) in control (data from Fig. 1h), neck-ablated and posterior-ablated pupae. Gray areas, ablated regions. **(c)** Ecad:3xmTagRFP notum live images upon LifeAct:GFP and PH:RFP670 clonal expression, and close-up (orange box). Yellow arrow, front protrusions; red arrows, cell rear devoid of protrusions. **(d)** Graph of the protrusion polarity AP component (mean ± SEM) as a function of the normalized AP position (*x*). Red arrows, polarity vectors; gray line, 0 of AP polarity. **(e)** Ecad:3xGFP and LifeAct:Ruby notum live images at 24 hAPF before (left) and 10 s after (right) AJ ablation. The ablated regions (orange dashed lines) are located either anteriorly (top) or posteriorly (bottom) to the LifeAct:Ruby-labeled clones. **(f)** Plot of the percentage of LifeAct:Ruby protrusion stretching (mean ± SEM), normalized to AJ recoil, upon anterior or posterior ablations in Ecad:3xGFP tissue at 24 hAPF. *** *p*=6.46 × 10^−7^ (Mann-Whitney U). **(g)** Ecad:3xmKate2 notum live images showing Dpy:YFP (top) and Sr:GFP (bottom) intensities. Dpy:YFP and Sr:GFP are projected above the AJ and at the level of the nucleus, respectively. **(h)** Graph of the mean rescaled velocity 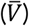 as a function of the normalized AP position (*x*) in control and *dpy*^*RNAi*^ tissues. In both control and *dpy*^*RNAi*^ dorsal-longitudinal indirect flight muscles (IFM) were ablated (see Extended Data Fig. 2h and Methods). **(i)** Graph of the protrusion polarity AP component (mean ± SEM) as a function of the normalized AP position (*x*) in control and *dpy*^*RNAi*^ tissues. Gray line, 0 of AP polarity. **(j)** Ecad:3xmKate2 and Myr:GFP notum live images before (left) and 10 s after (right) AJ ablations in control (top) and *dpy*^*RNAi*^ (bottom) tissues at 24 hAPF. The ablated regions (orange dashed lines) are located anteriorly to the Myr:GFP-labeled clones. **(k)** Plot of the percentage of Myr:GFP protrusion stretching (mean ± SEM), normalized to AJ recoil, upon anterior ablations in Ecad:3xmKate2 control and *dpy*^*RNAi*^ tissues at 24 hAPF. * *p*= 0.021 (Mann-Whitney U). **(l)** Graph of the mean rescaled velocity 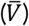 as a function of the normalized AP position (*x*) in control (data from Fig. 1h) and *sr*^*1*^ tissues. **(m)** Graph of the mean normalized Dpy:YFP intensity as a function of the normalized AP position (*x*) in control, *sr*^*1*^, and *dsh*^*1*^ tissues. Gray line, 0 of Dpy:YFP intensity. **(n)** Graph of the protrusion polarity AP component (mean ± SEM) as a function of the normalized AP position (*x*) in control (data from D), *sr*^*1*^, *dsh*^*1*^, and *sr*^*1*^*dsh*^*1*^ tissues. Gray line, 0 of AP polarity. **(o)** Graph of the mean rescaled velocity 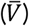 as a function of the normalized AP position (*x*) in control (data from Fig. 1h), *sr*^*1*^, *dsh*^*1*^, and *sr*^*1*^*dsh*^*1*^ tissues. Scale bars: 50 µm (a, g), 10 µm (c), 5 µm (e, j).

In the absence of anterior pulling or posterior pushing forces from surrounding tissues, we hypothesized that tissue flow is driven by the active migration of thoracic cells. The monolayered thoracic epithelium contacts both an apical extracellular matrix (aECM) and a basal extracellular matrix (bECM)^58,65–68^. During tissue flow, anterior notum cells exhibit a notable tilt, with their apical domains positioned more anteriorly than their basal counterparts (Extended Data Fig. 2a), suggesting that the aECM may serve as a substrate for anterior-directed migration. This is consistent with observations of collective flows, in which cells interact with the aECM or the vitelline membrane, during other morphogenetic movements in invertebrates and vertebrates^69–76^. Mosaic labeling of F-Actin with LifeAct:GFP and cell membranes using PH:RFP670 revealed the formation of F-Actin-rich apical protrusions, specifically at the anterior apical cell domain (Fig. 2c). Furthermore, high-resolution live imaging revealed F-Actin retrograde flows within these front protrusions (Extended Data Fig. 2b and Supplementary Video 5). Notably, quantification of protrusion polarity (Extended Data Fig. 2c) confirmed that protrusions are preferentially oriented towards the anterior (Fig. 2d). In addition, protrusion polarity is highest between 24 and 28 hAPF and in the anterior of the tissue, where the flow is faster (Fig. 2d and Extended Data Fig. 2d). To further test whether cells migrate on the aECM, we determined whether the protrusions attached to it. First, using ChtVis:Tomato as a live reporter of chitin in the aECM^68^, we found that the apical cell domain is in close contact with the aECM (Extended Data Fig. 2e). Second, we designed a live-imaging assay based on tissue recoil after multiphoton ablation, specifically severing the AJ to test protrusion attachment to the aECM (Extended Data Fig. 2f). In Ecad:3xGFP tissue with mosaic LifeAct:Ruby labeling, we ablated the AJ of cells located anterior to the front protrusions of LifeAct:Ruby labeled cells and recorded both Ecad:3xGFP and LifeAct:Ruby signals. Ecad:3xGFP revealed that the cells recoiled posteriorly, whereas LifeAct:Ruby showed that the front protrusions stretched as the cells recoiled (Fig. 2e,f). This suggests that the front protrusions are attached to the aECM. Furthermore, this attachment is specific to the front of the cell, as AJ ablation of cells posterior to the LifeAct:Ruby-labeled cells caused anterior recoil with minimal stretching of the LifeAct:Ruby-labeled apical cell domain at the rear (Fig. 2e,f). Collectively, these data indicate that the apical surface of anterior thoracic cells adheres to the aECM specifically at their front, consistent with the possibility that these cells leverage planar polarized apical protrusions for migration on the aECM.

To further establish that tissue flow is associated with migration and to understand its scaling with tissue size, we next aimed to delineate how cells interact with the aECM. The ZP-domain protein Dumpy (Dpy) mediates adhesion between the tissue and the aECM in *Drosophila*^58,77–79^. In the anterior notum, Dpy is produced by epithelial cells fated to become tendon cells^65^. As previously reported^58,65^, we found that Dpy:YFP is present between the tissue and the chitin during anterior thorax flows (Extended Data Fig. 2e). Interestingly, Dpy:YFP levels increased as cells started to migrate, consistent with a possible role in thorax cell displacement (Fig. 2g and Extended Data Fig. 2g). To evaluate the role of Dpy in thorax flow, we downregulated Dpy function through RNAi (*Dpy*^*RNAi*^) during tissue flow, and observed a substantial reduction in the anterior tissue flow (Fig. 2h and Extended Data Fig. 2h). Since *Dpy*^*RNAi*^ did not affect protrusion polarity (Fig. 2i), we hypothesized that Dpy instead controls attachment of the front protrusions to the aECM. To test this, we compared front protrusion stretching in control and *Dpy*^*RNAi*^ tissues after laser ablation. Consistent with a function in adhering cells to the aECM, *Dpy*^*RNAi*^ reduced front protrusion stretching upon cell recoil (Fig. 2j,k). Dpy is secreted by the epithelial tendon cells specified by the transcription factor Stripe (Sr)^80,81^. Accordingly, in the anterior notum, Sr:GFP localizes in a distinctive stripe pattern, each stripe presenting a noticeable AP gradient, mirroring the Dpy:YFP pattern (Fig. 2g and Extended Data Fig. 2g). Importantly, loss of Sr function using the *sr*^*1*^ allele diminished both the rescaled flow velocity and Dpy:YFP levels (Fig. 2l,m). We therefore conclude that, under the control of Sr, the aECM-interacting protein Dpy is required for anterior tissue displacement, reinforcing the idea that thorax tissue flow results from collective cell migration on the aECM.

Lastly, we sought to identify factors that could control protrusion polarity to analyze its contribution to tissue flow. In addition to the reduced flow velocity, the absence of Sr function diminished protrusion polarity (Fig. 2n). Given its role in planar cell polarization along the AP axis of the thorax tissue^82^, the Fz/Dsh planar cell polarity (PCP) pathway emerged as a possible regulator of protrusion polarity. In animals mutant for *dsh*^*1*^, a *dsh* allele that abrogates its PCP function^83^, both protrusion polarity and flow velocity were reduced without substantially affecting Dpy:YFP levels, corroborating the importance of protrusion polarity in tissue flow (Fig. 2m-o). Moreover, simultaneous impairment of Dsh PCP and Sr functions caused a further reduction in protrusion polarity and flow velocity compared with the single mutants (Fig. 2n,o). Together, these results indicate that both aECM attachment via Dpy and planar cell-polarized protrusions contribute to thorax tissue flow, confirming the existence of a migratory process on the aECM.

### Scaling of the morphogenetic field and modeling of collective cell migration

Having identified regulators of collective cell migration, we next examined how migration velocity scales with tissue size. Since migration depends on Sr expression, Dpy levels, and polarized cell protrusions (Fig. 2h,i and l-o), we analyzed whether these three quantities vary with animal size. As tissue length increased with animal size, the Sr:GFP and Dpy:YFP expression domains also lengthened, while the amplitudes of Sr and Dpy levels remained similar (Fig. 3a-c). Thus, gene patterning scales in length, as observed in multiple instances^1,2,84^. Furthermore, the polarity of cell protrusions did not vary with tissue length (Fig. 3d,e). These results agree with the fact that variations in tissue size were mainly due to changes in cell number, whereas individual cell apical size remained largely similar^60^ (Extended Data Fig. 3). We conclude that variations in Dpy levels, Sr patterning, and protrusion polarity along the AP axis scale with tissue length while keeping the same amplitude (Fig. 3f). We then aimed to determine whether their scaling with tissue length is sufficient to explain the experimentally observed scaling of migration velocity.

**Fig. 3.**
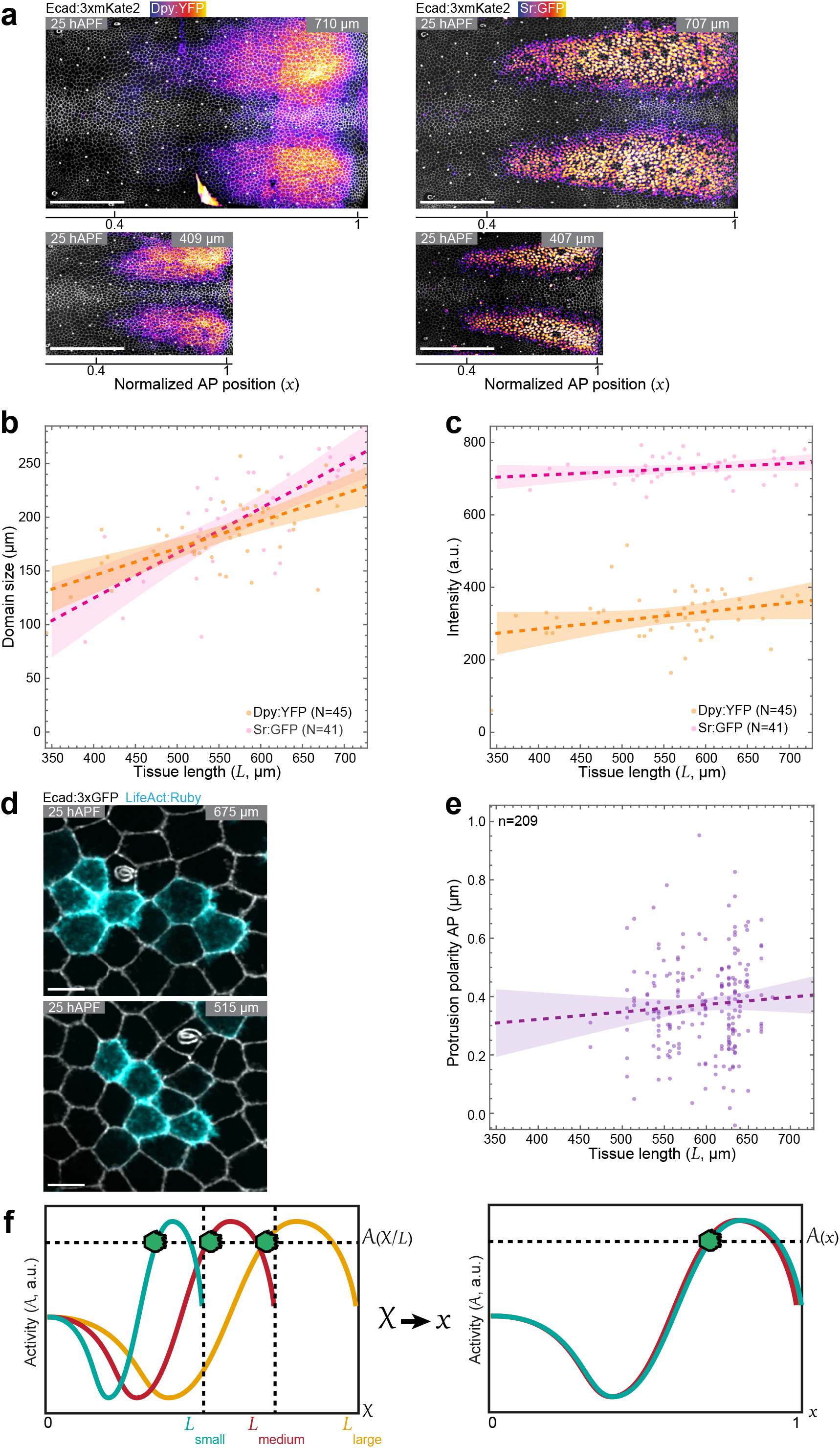
Morphogenetic field scales in length but not in amplitude with tissue length. The AP axis is horizontal with the posterior to the left. When relevant, the normalized AP position (*x*) is indicated below the images. Images were acquired in the anterior notum region at 25h APF. Quantifications are averaged from 24 to 28 hAPF. *p* value, Student’s *t*-test. Dashed line, linear fit; shaded area, 95% CI of the mean. N, number of animals. n, number of clones. Genotypes are listed in Extended Data Table 3. *x* = *X*⁄*L*. **(a)** Ecad:3xmKate2 notum live images showing Dpy:YFP (left) and Sr:GFP (right) intensities in large (top left, *L* = 710 µm; top right, *L* = 707 µm) and small (bottom left, *L* = 409 µm; bottom right, *L* = 407 µm) tissues. Dpy:YFP and Sr:GFP are projected above the AJ and at the level of the nucleus, respectively. **(b)** Graph of the Dpy:YFP and Sr:GFP domains sizes as a function of tissue length (*L*). Dpy:YFP, R^2^= 0.397, *p*= 3.51 × 10^−6^; Sr:GFP, R^2^= 0.496, *p*= 1.13 x10^−6^. **(c)** Graph of Dpy:YFP and Sr:GFP intensities as a function of tissue length (*L*). Dpy:YFP and Sr:GFP levels were averaged from 24 and 28 hAPF, and between *x* = 0.88 and 0.97 as in Fig. 1c. Sr:GFP, R^2^= 0.060, *p*= 0.12; Dpy:YFP, R^2^= 0.071, *p*= 0.077. **(d)** Ecad:3xGFP and LifeAct:Ruby notum live images upon LifeAct:Ruby clonal expression in a large (*L* = 675 µm) and a small (*L* = 515 µm) tissue. **(e)** Graph of the protrusion polarity AP component in the anterior notum (*x* = 0.88 to 0.97) as a function of tissue length (*L*). R^2^= 0.006, *p*= 0.28. **(f)** Schematic illustrating the inferred active force driving notum collective cell migration, expressed as a dimensionless activity (*A*), defined in the model based on Dpy levels and cell protrusion polarity, as a function of the AP position (left, *X*) or normalized AP position (right, *x*) in small, medium, and large tissues. After normalization of the AP axis, the activity profiles overlap, reflecting that the amplitudes of cell protrusion polarity and Dpy levels are similar in tissues of different lengths. Accordingly, the activity propelling cell migration at a given relative position (*x*) along the normalized AP axis is identical in small, medium, and large tissues. Scale bars: 100 µm (a, b), 5 µm (d).

Towards this goal, we developed a simple theoretical model of cell migration to determine the minimal ingredients for the scaling of flow velocity with tissue length to emerge around 25 hAPF (Fig. 4a, theory note and Extended Data Table 1). Since the tissue thickness (~10 µm) is much smaller than its AP length (>350 µm), and the flow is predominantly along the AP axis, we only considered the velocity along this axis (*V)* and wrote the model in one dimension (1D). We modelled the tissue as an active viscous material characterized by a viscosity coefficient *η*, as we were interested in the dynamics of the tissue at long timescales (hours, see theory note § 2.1). When the tissue deforms, each element of tissue experiences a viscous force (per unit length of material): 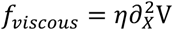. As the tissue migrates it also experiences friction with the ECM which leads to a friction force that we wrote as *f*_*friction*_ = −*ζV*, where *ζ* is the friction coefficient (Fig. 4a). Lastly, based on our experimental data (Fig. 2), we wrote the active force driving cell migration as depending on both Dpy levels and protrusion polarity. Since the AP profiles of Dpy levels and protrusion polarity scale with tissue length while maintaining the same amplitude (Fig. 3), we wrote the active force pattern as scaling with tissue length (while keeping the same amplitude) and depending only on the rescaled position along the AP axis, *x = X*/*L* (Fig. 3f). We therefore expressed the active force as proportional to a dimensionless activity term *A*(*x*) that combines the measured Dpy levels and protrusion polarity profiles along the AP axis, and a homogeneous activity (*A*_0_) accounting for the residual migration observed in *Dpy*^*RNAi*^ or *dsh*^*1*^*sr*^*1*^ double mutant conditions (Fig. 2h,l,o, 4b, Extended Data Fig. 4a-c and theory note § 2.3, 2.8, 2.9). The active force then reads: *f*_*active*_ = *γA*(*x*), where *γ* is a traction force per unit length. Writing the force balance on an elementary piece of tissue (of a given size, regardless of tissue length) and using the rescaled *x* position along the AP axis leads to the following equation of motion for the rescaled velocity 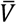:

**Fig. 4.**
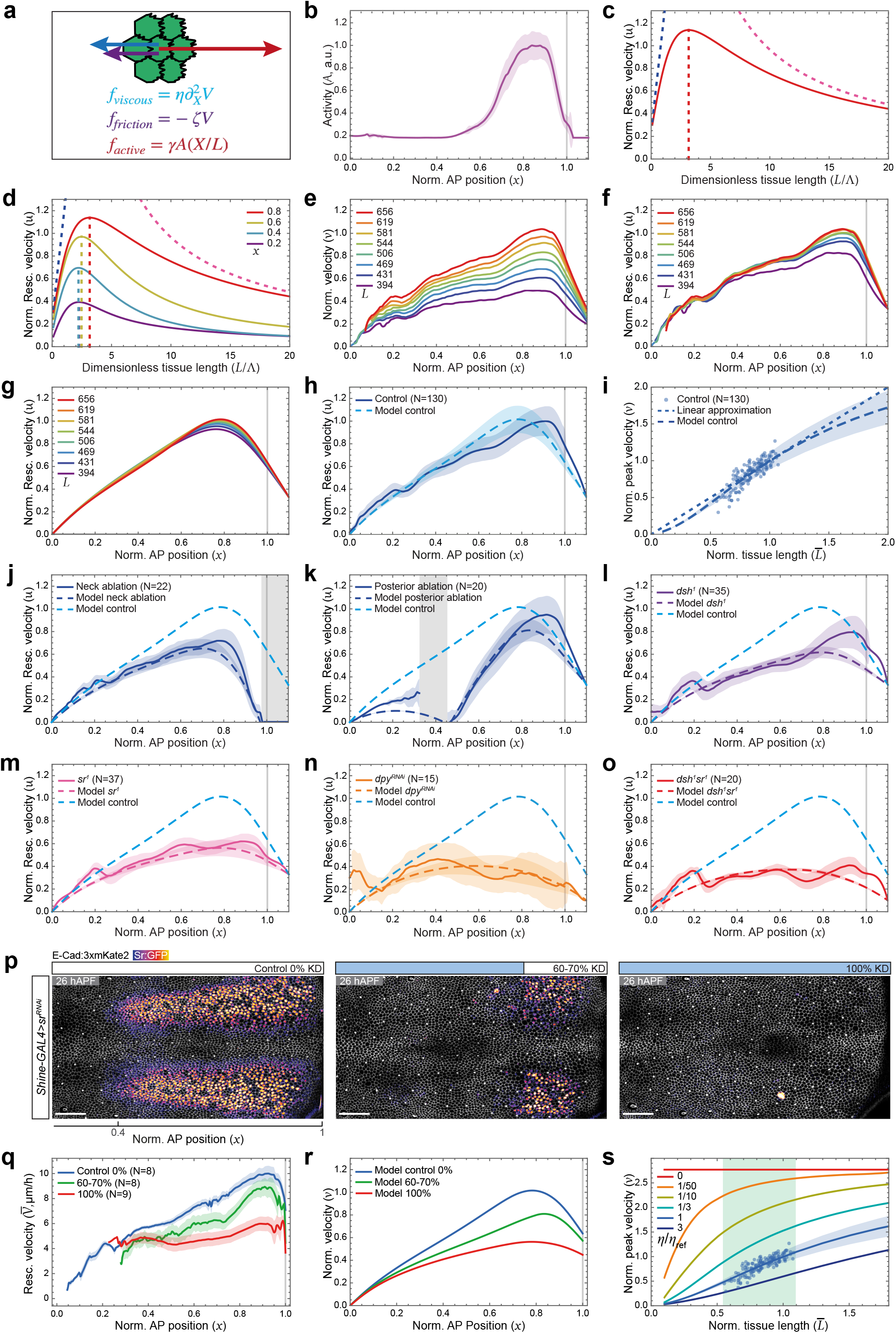
Minimal model of collective migration defines the contribution of tissue mechanical properties for scaling. The AP axis is horizontal with the posterior to the left. When relevant, the normalized AP position (*x*) is indicated below the images. Quantifications are averaged from 24 to 28 hAPF. Unless otherwise stated: solid line, mean experimental data; dashed line, model; shaded region, SD; vertical gray line (*x* >1), anterior limit of data (beyond this point data is extrapolated). N, number of animals. Genotypes are listed in Extended Data Table 3. 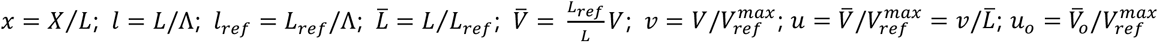. **(a)** Sketch of the 3 forces acting on a piece of tissue included in our modelling: a force due to tissue viscosity (*f*_*viscous*_), a friction force stemming from cell attachment to ECM (*f*_*friction*_), and an active force generated by cell traction on the aECM (*f*_*active*_). **(b)** Graph of the activity (*A*) as a function of the normalized AP position (*x*), defined by the AP profiles of Dpy levels and cell polarity, see also Fig. 3f and Extended Data Fig. 4a-c. **(c)** Graph of the model prediction for the normalized rescaled velocity (*u*) as a function of the dimensionless tissue length (*l*), at *x* = 0.8. Blue dashed blue line, viscosity dominated regime; pink dashed line, friction dominated regime; red solid red line, actual solution; red dashed line, maximum of actual solution. In the vicinity of this maximum *u* hardly depends on *l*, corresponding to a linear scaling of velocity. **(d)** Graph of the model prediction for the normalized rescaled velocity (*u*) as a function of the dimensionless tissue length (*l*) for different normalized AP positions (*x*) with *u*_*o*_ = 10. Blue and pink dashed lines, limit cases of dominating viscosity and friction, respectively (at *x* = 0.8, see C); vertical dashed lines, dimensionless tissue length corresponding to each curve maximum (*l*^∗^(*x*)). **(e)** Graph of the normalized velocity profiles (*v*) as a function of the normalized AP position (*x*) for tissue length bins (*L* = mean tissue length of the bin). Same data as Fig. 1h. **(f)** Graph of the normalized rescaled velocity (*u*) as a function of the normalized AP position (*x*) for different tissue length bins (*L* = mean tissue length of the bin). Same data as Fig. 1h. **(g)** Graph of the model prediction for the normalized rescaled velocity (*u*) as a function of the normalized AP position (*x*) for different tissue lengths (*L*) with *u*_*o*_ = 8.6. The Λ value is set so that *l*_*ref*_ = 3, thereby matching the superposition of the curves in experiments for the physiological tissue length range. **(h)** Graph of the normalized rescaled velocity (*u*) as a function of the normalized AP position (*x*) in control tissues and the associated model prediction. Shaded region in model, *A*(*x*) ± *σ*_*A*(*x*)_, where *σ*_*A*(*x*)_ is the standard deviation around *A*(*x*). Same data as Fig. 1h. **(i)** Graph of the predicted normalized velocity (*v*) as a function of the normalized tissue length 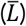. Data point velocities and tissue lengths were rescaled by the mean velocity in tissues of commensurate lengths (*V* = 9.7 μm/h for 616< *L* <707 µm) and the reference tissue length *L*_*ref*_ = 656 μm. Dashed straight line, linear approximation of the model curve at point 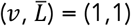, given by 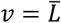. Same data as Fig. 1h. **(j)** Graph of the normalized rescaled velocity (*u*) as a function of the normalized AP position (*x*) in neckablated pupae and the associated model prediction (zero flow boundary condition at *x* = 0.97). Light blue dashed line, model prediction of control (see H); gray area, ablated region. Same data as Fig. 2b. **(k)** Graph of the normalized rescaled velocity (*u*) as a function of the normalized AP position (*x*) in posterior-ablated pupae and the associated model prediction (zero flow boundary condition at *x* = 0.45). Light blue dashed line, model prediction of control (see H); gray area, ablated region. Same data as Fig. 2b. **(l-o)** Graphs of the normalized rescaled velocity (*u*) as a function of the normalized AP position (*x*) in *dsh*^*1*^ (l), *sr*^*1*^ (m), *dpy*^*RNAi*^ (n) and *dsh*^*1*^*sr*^*1*^ (o) tissues with the associated model prediction. Dpy intensity SD was taken from the control condition in Fig. 2m, for *dpy*^*RNAi*^ and *dsh*^*1*^*sr*^*1*^ (see theory note § 2.11). Light blue dashed line, model prediction of control (see h). Same data as Fig. 2h, 2l, and 2o. **(p)** Ecad:3xmKate2 notum live images showing Sr:GFP intensities upon *Shine-GAL4>sr*^*RNAi*^ reduction of the Sr:GFP expression domain. Blue boxed activation regions; 0%, 60-70%, 100% reduction of the Sr:GFP domain length. **(q**,**r)** Graph of the rescaled AP velocity 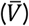 as a function of the normalized AP position (*x*) for different Sr:GFP domain lengths in *Shine-GAL4>sr*^*RNAi*^ tissues (q) and the corresponding model prediction (non-induced region, *A*(*x*)= *A*_*control*_(*x*); induced region, 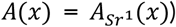 for the normalized velocity (*v*). Shaded area, SEM. **(s)** Graph of the model prediction for the normalized velocity (*v*) as a function of the normalized tissue length 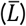 for different values of viscosity *η*/*η*_*ref*_. Light green vertical strip, range of physiological tissue lengths, including linear domain of the *η*/*η*_*ref*_ = 1 curve (located in the vicinity of 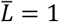. Lowering *η*/*η*_*ref*_ shifts the linear domain of the curves towards lower 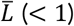, away from the physiological tissue length range (linearity of the scaling 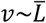 is gradually lost and gets closer to *v*~*cst*). Increasing *η*/*η*_*ref*_ shifts the linear domain of the curves towards higher 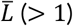, away from the physiological tissue length range (linearity of the scaling 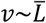 is gradually lost and gets closer to 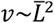. Same data as Fig. 1h; *η*/*η*_*ref*_ = 1 curve is replotted from (i). Scale bars: 50 µm (p).

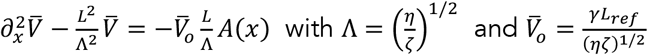

In addition to *A*_0_, this model therefore has two adjustable parameters, 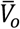 and Λ. 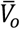 is a characteristic rescaled velocity that determines the amplitude of the migration velocity. Λ is the characteristic hydrodynamic length, defined by two tissue mechanical properties, viscosity, and friction with the ECM. This length determines the length scale over which a mechanical perturbation propagates within the tissue. Since Λ determines the hydrodynamic behavior of the tissue, it is not the absolute tissue length *L* but rather the ratio *L*/Λ, namely the dimensionless tissue length, that appears in the equation.

### Minimal model of migration reproduces scaling of velocity and supports a key role for tissue mechanical properties

This minimal 1D model led to several important findings. In the case of linear scaling of *V* with tissue length *L*, the rescaled velocity 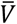 (proportional to *V* divided by *L*) is therefore independent of *L*. Interestingly, the tissue length *L* explicitly appears in the equation of motion, making the experimental independence of 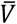 with *L* (or equivalently, the experimental linear scaling of *V* with *L*) non-trivial, e.g. not solely depending on the scaling of the morphogenetic field length (modelled by *A*(*x*) in the equation). In the general case, this equation shows that 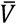 depends on *L*, and the dependence of *V* on *L* is nonlinear (see theory note § 2.2-2.5). Therefore, according to this model, the observed scaling of migration with *L* cannot arise for all tissue lengths. To understand how a linear scaling can occur, we first analyzed two limit cases of the model (Fig. 4c). In the limit case where the tissue length is much larger than the tissue’s hydrodynamic length scale (*L* ≫ Λ, i.e., friction dominates), 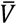 decreases with *L* with a 1/*L* dependence (i.e., *V* is constant for all tissue lengths). In the other limit case, namely *L* ≪ Λ (i.e., viscosity dominates), 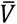 now increases with *L* with a linear dependence (i.e., *V* scales in *L*^2^). From the analysis of the main equation and of these two limit cases, we conclude that: (i) the relevant variable to explore the scaling behavior is the ratio of the tissue length and its hydrodynamic length scale (*l* = *L*/Λ); (ii) for a given range of tissue lengths, the scaling behavior of *V* therefore depends on the tissue mechanical properties; (iii) between these two limit cases, for a given range of tissue lengths, the general solution 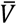 must admit a maximum set by the tissue mechanical properties (Fig. 4c); 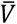 is then independent of *L* in the vicinity of this maximum, which implies that *V* varies linearly with *L*, as observed experimentally. Therefore, for a given range of tissue lengths, linear scaling of migration velocity with tissue length emerges when tissue mechanical properties are properly tuned (Fig. 4c). More generally, this illustrates that tissue mechanics can determine the scaling behavior of velocity with tissue length.

To further confirm this result, we analyzed the solutions of the equation for 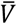 (see theory note § 2.7-2.9). We first explored these solutions by representing 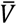 along the AP axis for different values of the dimensionless tissue length *L*/Λ (Extended Data Fig. 4d), and as a function of *L*/Λ for different positions *x* (Fig. 4d). We found that 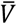 indeed displays a series of maxima when plotted against *L*/Λ for different *x*, which differ but remain fairly close to one another (2.2<*L*/Λ<3.4, Fig. 4d, Extended Data Fig. 4e,f and theory note § 2.9). Hence, we do not expect migration velocity to scale perfectly along the entire AP axis, as a given range of tissue lengths cannot be centered simultaneously on 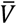 maxima at both the rear and the front of the tissue (see theory note § 1.2-1.5, 2.9). We therefore analyzed in more detail the *V* and 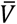 AP profiles for different tissue length bins and confirmed this prediction experimentally: the rescaled velocity AP profiles collapse for 0<*x*<0.7, indicating nearly perfect linear scaling with *L*, but remain slightly separated for *x*>0.7 (Fig. 4e,f). To then determine whether the model could reproduce the observed scaling, we set the characteristic length Λ, or more accurately the ratio *L*/Λ, so that the physiological range of tissue lengths is centered on the range of maxima of the 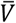 versus *L* curves corresponding to the rear part of the tissue (0<*x*<0.7) (Extended Data Fig. 4g-j and theory note § 2.9). This therefore leads to the superposition of the model rescaled velocity curves of each tissue length bin, with more discrepancy near the neck, as observed experimentally (Fig. 4e,f,g). We found Λ to be about 220 µm and then confirmed that the model describes the control data, specifically the normalized rescaled velocity profile and the seemingly linear *V*(*L*) relationship (Fig. 4h,i). Thus, in the physiological tissue length range, this minimal model successfully reproduces the observed scaling of migration velocity with tissue length.

### The theoretical model can account for the different experimental perturbations

With all three adjustable model parameters now set, we performed three independent tests to validate the model. First, we compared its predicted rescaled velocity profiles for laser ablation experiments in which we cauterized the tissue at the anterior or posterior domains (Fig. 2a,b). By mechanically perturbing the tissue at specific locations, these experiments probe the characteristic length scale Λ that determines tissue hydrodynamic behavior. We found a fair agreement between our prediction and the measured migration velocities in both experiments, thereby showing that our estimation of Λ is correct (Fig. 4j,k). Second, we compared the model predictions with mutant conditions whereby we disrupted tissue PCP (*dsh*^*1*^), Dpy level (*Dpy*^*RNAi*^ or *sr*^*1*^), or both (*dsh*^*1*^*sr*^*1*^). To do so, for each mutant condition, we modified the activity function *A*(*x*) according to Dpy levels and protrusion polarity profiles measured experimentally in each condition (Extended Data Fig. 4k). Again, as observed experimentally, the model predicts a decreasing migration velocity matching the experimental data (Fig. 4l-o). Last, we analyzed the impact of reducing the length of the Sr domain in the model. The model predicts that a drastic reduction in the length of this domain triggers a substantial decrease in velocity. To test this prediction experimentally, we aimed to mimic a 70% reduction in the length of the Sr domain along the AP axis. For this purpose, we used the optogenetic Shine-GAL4 system^85^ to specifically express *sr*^*RNAi*^, thereby reducing the Sr domain length in large animals (Fig. 4p). Strikingly, as anticipated by the model, the migration velocity decreased in this experimental condition (Fig. 4q,r). Altogether, these results show that very simple assumptions about tissue rheology and activity are sufficient for the scaling of migration velocity with animal size to emerge. This phenomenon is nevertheless non-trivial as it predicts that it only occurs within a given crossover regime between viscosity and friction dominated flows and therefore cannot solely result from scaling of the morphogenetic field.

### Modulation of cell-cell adhesion and cell contractility changes the scaling behavior of the tissue with animal size

Lastly, we investigated the contributions of tissue mechanical properties to the AP velocity profile and its scaling behavior. According to the model, within the physiological range of animal lengths, lowering viscosity should make the migration velocity *V* less dependent on tissue length, thereby altering its scaling (Fig. 4s, Extended Data Fig. 4l-o and theory note § 2.12). Previous reports have established that lowering either cell-cell adhesion or cell contractility can decrease epithelial tissue viscosity^86–90^. In particular, loss of p120-catenin (p120) was shown to reduce tissue viscosity by enabling larger cell deformation due to an increase in Ecad turnover^91^. We recorded tissue flow in *p120* mutant tissues from animals of different sizes and found that the dependence of migration velocity on tissue length was indeed decreased relative to control tissues (Fig. 5a). Importantly, we verified that tissue contraction-elongation was larger, and cells deformed more in the absence of p120 function (Extended Data Fig. 5a), consistent with a reduction in viscosity^91^. We then explored whether a reduction in tissue viscosity in the model was sufficient to account for the observed migration velocity profile and altered scaling between velocity and tissue length in *p120* animals. To this end, we first measured the Dpy:YFP intensity profile and protrusion polarity upon loss of p120 function. While the Dpy:YFP profile was not affected, protrusion polarity was reduced by around 50% (Extended Data Fig. 5b,c). Using these experimental values, we modified the *A*(*x*) profile accordingly (Extended Data Fig. 5d) and could recapitulate the *p120* migration velocity profile and its scaling behavior by setting *p120* tissue viscosity to about 1/5 of the control condition (Fig. 5b,c and Extended Data Fig. 5e,f). With this reduction, the model predicts minor changes in the AP velocity profile, a prediction that was experimentally observed (Fig. 5b and Extended Data Fig. 5g-i). To further confirm the impact of tissue viscosity on migration velocity scaling, we reduced it through a different mechanism, namely by lowering Myosin II-dependent contractility^87^. We found that the expression of a dominant negative form of non-muscle Myosin II heavy chain (*zip*^*DN*^) led to a reduced scaling of migration velocity as a function of tissue length, showing the impact of cell contractility on the scaling properties of the tissue (Fig. 5d). The reduction of cell contractility also led to a major change in the AP velocity profile (Fig. 5e and Extended Data Fig. 5j), with a steeper velocity slope in the anterior tissue. Since cell contractility is known to play an important role in force generation during migration^26–37,92^, we first reduced the active force in the model (*γ*) to explore the agreement between the experimental data and the model. Both the scaling behavior and the steeper slope along the AP axis could then be recapitulated in the model by reducing the viscosity by a factor of about 1/10 (Fig. 5e,f and Extended Data Fig. 5k-n). Together, these experimental data indicate that lowering tissue viscosity, through the reduction of either cell-cell adhesion or cell contractility, alters the scaling properties of the tissue within the physiological range of animal sizes. Combining our experimental and theoretical findings, we propose that mechanical properties play a critical role in the scaling of collective cell migration with animal size.

**Fig. 5.**
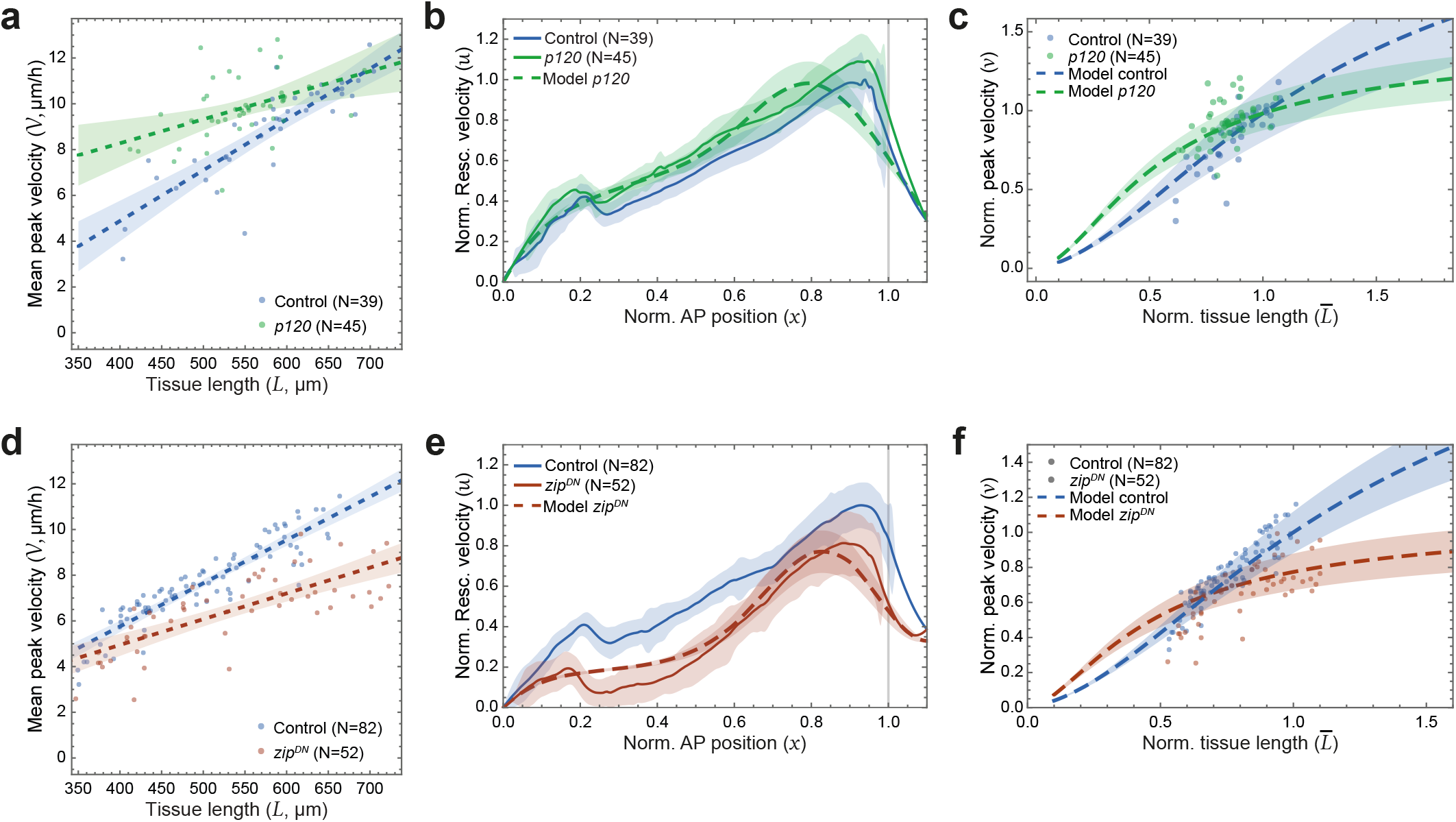
Cell-cell adhesion and cell contractility impacts scaling of velocity amplitude with tissue length. The AP axis is horizontal with the posterior to the left. Quantifications are averaged from 24 to 28 hAPF. Unless otherwise stated: solid line, mean experimental data; dashed line, model; shaded region, SD. Vertical gray line (*x* =1), anterior limit of data (beyond this point data is extrapolated). N, number of animals. Genotypes are listed in Extended Data Table 3. Interactions between genotype and size (A and D) are tested with ANOVA2. 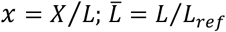. **(a)** Graph of the mean peak velocity (*V*) as a function of tissue length (*L*) in control and *p120* tissues. Dashed lines, linear fits. Shaded area, 95% CI of the mean. *p*= 3.25 × 10^−3^. **(b)** Graph of normalized rescaled velocity (*u*) as a function of the normalized AP position (*x*) in control and *p120* tissues of commensurate lengths (622 µm < *L* < 704 µm). For the model prediction, cell polarity and Dpy levels in *p120* tissue were set with experimental measurements (see Extended Data Fig. 5b,c). The model *u* profile was fit by only adjusting the tissue viscosity *η* to recover the *u* experimental maximum value (*η*_*p*120_/*η*_*control*_ = 0.20). **(c)** Graph of the predicted normalized velocity (*v*) as a function of the normalized tissue length 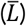 in control and *p120* tissues. Data point (same as A) velocities and tissue lengths were rescaled by the mean velocity in tissues of commensurate lengths (*V* = 10.6 μm/h for animal length range: 623 µm < *L* < 704 µm). **(d)** Graph of the mean AP velocity (*V*) as a function of tissue length (*L*) in control and *zip*^*DN*^ tissues. Dashed lines, linear fits. Shaded area, 95% CI of the mean. *p*= 3.06 × 10^−5^. **(e)** Graph of normalized rescaled velocity (*u*) as a function of the normalized AP position (*x*) in control and *zip*^*DN*^ tissues of commensurate lengths (615 µm < *L* < 713 µm). For the model prediction, cell polarity and Dpy levels in *zip*^*DN*^ tissue were taken similar to control, resulting in same A(*x*). The model prediction of *u*(*x*) was fit by adjusting cell traction force *γ* and tissue viscosity *η* to recover the *u* experimental profile shape and peak amplitude (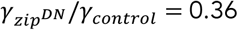 and 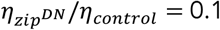). **(f)** Graph of the predicted normalized velocity (*v*) as a function of the normalized tissue length 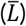 in control and *zip*^*DN*^ tissues. Data point (same as D) velocities and tissue lengths were rescaled by the mean velocity in tissues of commensurate lengths (*V* = 9.9 μm/h for animal length range: 615 µm < *L* < 663 µm).

## Discussion

Size is a fundamental attribute of any biological system. Our knowledge of tissue size control has vastly progressed in recent years^5–7^. While large size variations can be observed within or across species, how organisms adapt their tissue flow dynamics to size has remained largely unexplored. In this study, we investigated this fundamental question during collective cell migration and provided novel principles regarding the scaling of tissue dynamics with size. Leveraging experiments and theory, we found that collective migration adapts to tissue size through linear scaling of its velocity as a function of tissue size, a sufficient condition for correctly positioning cells independently of tissue size (since migration duration is constant across sizes). We uncovered that genetic patterning alone is insufficient to ensure proportional scaling of tissue velocity across all sizes. Instead, linear scaling emerges within a crossover regime of tissue mechanical properties tuned in relation to the physiological range of animal sizes. Our findings could have major consequences for developmental plasticity and robustness, regeneration, as well as implications for the constraints enforced on tissue dynamics and size during evolution.

Scaling with size is a core property of biological systems^93–96^. During development, this is highlighted by seminal experiments on embryo size reduction at the beginning of the last century^1^. The theoretical framework of morphogen gradients and its implications for cell fate specification have been instrumental to investigate the mechanisms of such developmental scaling^1^. A large body of work has delineated how cell fate patterning can become largely invariant to tissue size, producing similar proportions and distributions of cell fates independently of animal size. This can rely on the scaling of morphogen gradients, generated by distinct mechanisms mainly rooted in a length scale linked to the ratio between morphogen diffusion and degradation^1,2^. Studies have highlighted that linear scaling of gene patterning with size can be achieved in certain modalities of morphogen gradient scaling, and that gene regulatory networks also directly contribute to correct cell fate specification during tissue growth^1,2^. However, since tissue flow dynamics impose the production of mechanical forces to displace cells, it has remained unclear how the scaling of gene patterning can account for the adaptation of tissue dynamics to tissue size. In addition, varying embryoid body size *in vitro* has revealed that the scaling of gene patterns with gastruloid size does not equate to the scaling of tissue dynamics^97^, suggesting that, at least *in vitro*, distinct mechanisms account for their size invariance. Here, we found that the velocity of collective epithelial cell migration linearly scales with tissue length. This linear scaling occurs within a crossover regime of tissue mechanical properties that set the tissue hydrodynamic length scale, defined by the (square root of the) ratio of viscosity to friction. Importantly, given tissue rheological properties, linear scaling and robust cell allocation thus occur when these mechanical properties are appropriately tuned to the physiological range of animal sizes. Interestingly, our analysis also highlights that perfect scaling of velocity amplitude with animal size cannot be achieved along the whole tissue. Altogether, we conclude that the scaling of tissue development with size emerges from at least two distinct length scales, a first one related to the scaling of the morphogenetic field regulating tissue patterning, and a second one related to the length scale at which mechanical forces propagate within the tissue.

Tissue flows depend on force production and the tissue mechanical properties, including interactions with the ECM. The analysis of tissue dynamics has outlined how gene patterning can modulate force production via changes in the organization and dynamics of the cytoskeleton and in cell adhesion^3,98,99^. While tissue mechanical properties remain challenging to measure and manipulate in developing systems, recent studies have explored how properties such as elasticity, viscosity, and friction with the ECM contribute to tissue deformation^88,90,100^. Numerous studies have delineated how cell contractility and cell-cell adhesion determine tissue viscosity and stress dissipation during tissue shaping^86–88,90^. Our results highlight that the scaling of gene patterns ensures the increase of total tissue force production for larger tissue sizes, and that tissue mechanical properties set the scaling behavior of tissue flow with animal size. Together, these findings make mechanical properties a critical tissue attribute, defining not only the magnitude of tissue flow but also its adaptation to size.

Collective cell migration is a fundamental process in tissue development, homeostasis, and repair, and also plays a key role in cancer metastasis^21^. So far, most emphasis has been put on mechanisms driving migration on the basal ECM, where protrusive force is often generated by retrograde flows of actomyosin that transmit force to the ECM via integrins and propel the cell forward^23–25^. Notably, many tissues harbor an apical ECM with which they interact during development^69–76^. In this study, we aimed to achieve sufficient knowledge of the process of migration on the apical ECM to model and explore the scaling of collective migration with animal size. A major difference between migration on the apical versus basal ECM lies in the transmembrane molecules that transmit traction forces to the ECM^23–25^. Our theoretical model is agnostic to such molecular details and uses basic modeling ingredients whose relevance has been established for collective migration on basal ECM^28,31,43,46–53^. We therefore anticipate that our findings will advance the understanding of size-dependent scaling of collective cell migration in multiple contexts, both *in vivo* and *in vitro*.

Developmental timing varies across species and is influenced by gene expression dynamics, metabolism, and protein turnover^101–105^. Drastic changes in developmental duration are associated with large differences in size and shape observed across species^96^. Our work focused on size variation within the same species, and we found that the scaling of collective cell migration occurs without changes in its duration. Within a species, the temporal control of developmental processes is also regulated by systemic signals, metabolic rates, and hormones^6,106^. It will therefore be relevant to probe how, within a given species, collective flow duration can be made independent of animal size, even though the organs producing these systemic signals likely differ in size and exhibit distinct metabolic rates linked with variations in animal size.

Collective cell flows lie at the heart of development, homeostasis, and repair. Our work opens numerous avenues for research into the mechanisms by which other 2D and 3D tissue dynamics scale with tissue or organ size. More generally, scale effects appear in three different contexts^93^: (i) within the ontogenetic sequence of an organism; (ii) among individuals of a given species; and (iii) across species. On the one hand, this calls for exploring how the interplay between tissue gene patterning and mechanical properties contributes to the variation in tissue flow during development, both within and across species. On the other hand, it highlights the need to delineate how tissue mechanical properties are set, and how they influence the robustness and evolution of tissue flow adapted to animal size variations arising from intrinsic and environmental cues.

## Methods

### Fly stocks and genetics

Extended Data Table 2 presents a compilation of *Drosophila melanogaster* stocks that were utilized in this study, along with their corresponding references. Extended Data Table 3 provides the genotype used in each figure panel if applicable. *Drosophila* stocks and crosses were maintained on standard molasses/cornmeal/yeast food at 18 °C or 25 °C. Small pupae were obtained by transferring early instar larvae to tubes containing 7% agar before pupariation. Loss-of-function and gain-of-function experiments were conducted using the Gal4/UAS system, with the Act5C-GAL4 driver in conjunction or not with the temperature-sensitive Tub-Gal80^ts107,108^ to temporally regulate transgene expression. Thereto, animals were raised at 18 °C to prevent transgene expression. Subsequently, they were subjected to a temperature shift to 29 °C for a period of 30-34 hours to induce expression. The optogenetic Shine-Gal4 system^85^ was used to spatially modulate the *sr* expression domain. To generate small clones expressing UAS-LifeAct:Ruby, UAS-LifeAct:GFP, UAS-PH:RFP670 or UAS-Myr:GFP for protrusion polarity analysis and attachment analyses, we utilized the FLP/FRT flip-out system^109,110^. Thereto 0 hAPF pupae were heat-shocked for 5 min at 37 °C and imaged at the appropriate timing.

### Generation of Sr:GFP and PH:RFP670 transgenic lines

The functional *Sr:GFP* allele was generated through CRISPR/Cas9-mediated homologous recombination at the endogenous locus, utilizing the vas-Cas9 line^111^. Guide RNAs were inserted into the pCFD5:U6:3-t:gRNA vector^112^. The following guide RNAs were used: 5’-TGG TTT GCG GTA ZGA GGG CGT GG-3’ and 5’-TGG CGG TGG TTT GCG GTA GGA GG-3’. Homology sequences (HR) were cloned into homologous recombination vectors containing an hs-mini-white cassette flanked by two loxP sites^113,114^. Additionally, an N-terminal GFP sequence was inserted. The vector for GFP tagging has been previously described^113^. The following HR1 and HR2 sequences were cloned into this tagging vector: (HR1) 5’-CTA ATT ATG GGG TGT CGC CCT TCG GGT CTC TAG TTG AAC GAA GAG TTC TAT GGC ATT CCG-3’ and 5’-GAA CTG CCT GAA GAA CCG CTG GAC CCC GAA CTG GAG GGC GTG GCT GGC GAC TTG-3’; (HR2) 5’-CTC CGG AAG TGG TAG CTC AGG GTC TAG TGG ATA CCG CAA ACC ACC GCC ATA TG-3’ and 5’-GCC CTT GAA CTC GAT TGA CGC TCT TCG TCC GAA TTA TCC AAA GGG GAG CTT GTG-3’.

UAS-PH:RFP670 transgenic lines were generated by cloning a codon optimized RFP670 (synthetized by Integrated DNA Technologies), a SSGSSGSSGS linker and the Phospholipase C**δ**1 PH domain (as in ^115^) sequences using the following primers pUAS-RFP670: 5’-TTA CTT CAG AAC TTA AAA AAA AAA ATC AAA ATG GTA GCA GGT CAT GCC TCT G-3’, Linker-RFP670: 5’-CGA GCC ACT CGA TCC CGA ACT ACC CGA CGA GCT CTC AAG CGC GGT GAT CCG-3’, Linker-PH: 5’-ATC GCG ACG CGG ATC ACC GCG CTT GAG AGC TCG TCG GGT AGT TCG GGA TCG-3’ and PH-P10: 5’-AAT TGA TTT GTT ATT TTA AAA ACG ATT CAT CTA TTG CCG CTG GTC CAT GCT TC-3’ into the pUAS-IVS-SYN21-P10 transgenic vector (Addgene). Cloning was carried out using SLIC^116^. Embryo transgenesis was conducted by Bestgene, and transgenes were validated by sequencing. Detailed plasmid maps and DNA sequences are available upon request.

### Pupa mounting and imaging

Pupae were collected either at 0 hAPF or at head eversion (12 hAPF). The pupa mountings for live imaging were adapted from previous protocols^55,59^. At approximately 14-17 hAPF, the dorsal part of the pupal case covering the head and thorax was removed. Subsequently, the pupae were affixed ventrally onto a glass slide, heads were elevated using six layers of double-sided tape. Two coverslip spacers were positioned near the edges of the slide. The imaging coverslip was coated with a fine layer of mineral oil 10 S (VWR International) before mounting and sealing with nail polish. Pupae expressing a fluorescently tagged Ecad AJ marker, were, for tissue flow analyses imaged at either 25 °C or 29 °C, using a 40x/1.4 OIL DIC H/N2 PL FLUOR or 40x/1.3 OIL DIC H/N2 PL FLUOR objective on inverted spinning-disk confocal microscopes equipped with CSU-W1 unit (Yokogawa): (1) Andor/Roper/Nikon wide system with an sCMOS camera (Orca Flash4, Hamamatsu) and Borealis module (Andor), (2) Roper/Zeiss wide system with an sCMOS camera (Orca Flash4, Hamamatsu), a homogenizer (Visitron) and FRAP/ablation module (GATACA systems), (3) Roper/Nikon/GATACA wide system with an sCMOS camera (BSI camera, 95% QE) and homogenizer (Visitron), or (4) Roper/Zeiss/Spark wide system with an sCMOS camera (Orca Flash4, Hamamatsu) and homogenizer (Visitron). Multiple position images, typically 2 per tissue, were acquired at 15-min intervals (28-30 µm z-stack with 1 µm spacing), typically for 20-22 h. For imaging protrusion dynamics, pupae were imaged at 3 s intervals using a 63x/1.4 OIL DICII PL APO objective on an inverted laser scanning microscope (LSM880 NLO, Carl Zeiss) equipped with a ZEISS Airyscan detector module.

### Z-stack projection

2D projections were performed either with the Fiji^117^ plugin Local-Z-Projector^118^ or a previously published pipeline^119^ using both Fiji and MATLAB (https://www.mathworks.com/), subsequently multiple positions were stitched using Fiji’s default plugin “Image Stitching”^120^. In both cases the apical surface was determined based on the fluorescent Ecad signal. Briefly, in the case of our previously published pipeline the background was subtracted using MosaicSuite^121^, the images were then downscaled by a factor of 10, followed by the application of a variance filter along the z-direction. The resulting stack was then used by a MATLAB script to determine the most apical variance peak along the z axis for each xy position with *peakseek* (https://www.mathworks.com/matlabcentral/fileexchange/26581-peakseek). This process generated a topographic map (z-map) of the apical surface. The generated apical z-map was then used to project the fluorescent Ecad signal for tissue flow velocity, signals above the junctions for Dpy:YFP level measurements and apical protrusions polarity and stretching, or signals below the junctions for nuclear Sr:GFP quantifications.

### Quantifications of tissue dynamics

#### Spatial and temporal registration of time-lapse movies

Spatial alignment was performed using the neck-thorax (N-T) boundary^122^, the two anterior dorsocentral (aDC) macrochaetes and the midline. We then averaged signal from the two medial hemi-thoraces, defined by the midline and the lateral positions of the aDC macrochaetes; all measurements being expressed in reference to the right hemi-thorax. To determine the length of the notum, we first established on commensurate animals that the mean notum length is 1.75 times the distance between the N-T boundary and the mean AP position of the two aDC macrochaetes. Subsequently, the tissue length for each animal was determined by multiplying the distance along the AP axis between the N-T boundary and the mean position of the two aDC macrochaetes by 1.75 to obtain the full notum length. In all time-lapse movies, the midline was aligned with the horizontal *x* axis. The temporal alignment was performed in two-steps. First, the final division of the most anterior microchaetes was manually annotated at 22 hAPF. Secondly, when possible, we temporally aligned the anterior flows of each pupa so that they reached ¾ of their maximum velocity at the same time. The mean of the manually annotated values was then used to define 22 hAPF for all movies from the same condition. This second correction was not possible in *Dpy*^*RNAi*^ and *sr*^*1*^*dsh*^*1*^ tissues; in this case the temporal alignment was done using only the final division of the most ant erior microchaetes. To account for differences in developmental time between experiments conducted at 25 °C or 29°C, a correction factor of 1.27 was applied to data acquired at 29 °C, aligning it with the 25°C timing^123^. All experiments conducted at different temperatures were performed alongside matched controls.

#### Lagrangian tissue velocity measurements, rescaled velocity, and tissue deformations

Local tissue velocities were determined using particle image velocimetry (PIV) on projected stacks using a 5×5 µm^2^ grid^54^. We used a Lagrangian description of tissue flow through tracking of tissue sub-regions, based on their initial positions and following their evolution over time. This enables to follow the changes in the velocity of tissue patches according to their initial genetic identity and position. The “pseudo-tracking” was performed using local tissue velocities calculated through PIV to transform the Eulerian-based mapping of cell flows into Lagrangian maps. In short, we initially obtained Eulerian flow maps (*x, y, t*), applied masks to identify regions outside the tissue, and interpolated missing values. We then applied temporal (1.25 h) and spatial (7% along AP axis and 28% along ML axis) average smoothing on these datasets. Using these processed maps, we then performed pseudo-tracking of each tissue region to calculate velocities in a Lagrangian manner. The resulting velocity maps were smoothed in time (1.25 h).

To compare the cell shape changes in control and *p120* tissues, movies were segmented and tracked to determine tissue deformation (elongation-contraction along the AP axis) and the associated cell shape changes using the formalism that relates tissue deformation to cell dynamics as described in^54,55^. For this purpose, cell shape changes were estimated in tissue regions of reference size 90×90 µm^2^ that were adjusted to the size and aspect ratio of each animal to make sure each region was comparable between animals ^54^.

After averaging the two hemi-thoraces and temporal alignment, the kymographs were obtained by averaging the kymographs from different animals. To generate the velocity standard deviation kymographs, the velocity standard deviation among animals was calculated for each pixel in the kymograph. Pixels in the mean kymograph covered by fewer than 3 animals were filtered out. Similarly, rescaled velocity kymographs were generated using 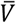 instead of *V* (see Extended Data Table 1).

For measurements of flow duration, the time interval between the two time-points for which the tissue flow velocity is above 20% of *V*_*max*_ was determined for each pupa.

### Quantification of apical protrusion polarity

To quantify protrusion polarity, high-resolution images of Ecad:3xGFP or Ecad:3xmKate2 (AJ signal) in tissues harboring LifeAct:Ruby or Myr:GFP (protrusion signal) somatic clones were projected. Isolated small clones were then manually selected to measure the polarity of their protrusion as follows (Extended Data Fig. 2c). For each clone, the AJ signal was automatically segmented and manually corrected. The protrusion signal was segmented with a manual threshold. Using a custom Fiji macro, each cell within a clone was divided into 24 angular sectors. For each sector, both the length of segmented AJ signal and the area of segmented protrusion signal extending beyond the segmented AJ signal were measured (up to 5 µm away from the AJ). The resulting area was then divided by the length of the AJ signal for each sector to obtain the normalized protrusion spread. Each segmented cell was dilated (by 10 pixels), and the overlap was used to identify sectors pointing towards other cells of the clone and exclude them. Identical angles within a clone were averaged resulting in 24 angular measurements per clone. The first harmonic of the circular Fourier decomposition was then extracted to define a polarity vector per clone. The orientation of the polarity vector represented the main direction of protrusion spreading, while the length of the vector reflected the amplitude of polarization. Polarity vectors were projected onto the AP axis to obtain a quantitative score of AP polarization for each clone. The coordinates of each clone within the thorax were calculated using an overview picture (not shown). The midline orientation was used to aligning the angular distribution of each clone with AP axis. Lagrangian coordinates for each clone were then inferred from the mean velocity kymographs of the corresponding genotypes.

### Quantification of cell area and density

Mean cell areas were determined by segmenting a strip of tissue located between *x* = 0.43 and *x* = 0.89 along the AP axis, and between 50% and 70% of the width of the ML axis (between the midline and one aDC macrochaetae, see Fig. 1b) in each animal using the same segmentation script used for protrusion polarity measurements. The rectangular region with the same AP length but spanning the whole ML axis width was used to estimate the cell number in the region by dividing this region area by the mean cell area initially determined from the segmented strip.

### Optogenetic manipulation of the Sr expression domain

Spatial modulation of the *Sr* expression domain was accomplished by using the light-activatable Shine-GAL4 system^85^. For these experiments, animals were reared in the dark, and large pupae were mounted under amber light conditions. Pupa were pre-incubated for 15-30 min at 29 °C prior to imaging at 29 °C to increase the efficacy of Magnets dimerization upon blue light exposure. In large (*L* > 700 µm) early pupa (12 hAPF) expressing the Ecad:3xmKate2 AJ marker as well as Sr:GFP, UAS-dsRNA expression targeting *sr* (*sr*^*RNAi*^) was induced upon single-photon 491 nm illumination (100 ms pulse at 0.72 mW) using an inverted spinning-disk microscope (Andor/Roper/Nikon) in binning 2 mode and using a 40x/1.3 OIL DIC H/N2 PL FLUOR objective. In each pupa the two bi-lateral tissue halves (covering the entire notum medial-lateral tissue axis) were activated, utilizing the tissue macrochaetes as spatial references. The ROI of activation (camera field of view, ~115600 µm^2^) was set as follows: pupae were placed on the microscope holder along the animal AP axis (anterior to the left), where the top or bottom ROI references were placed on the midline, while the right reference reflects the positioning of the activation ROI along the AP axis: when placed between the pDC and aDC, or between the pDC and aSC, this yielded a reduction of the *sr* expression gradient by 100%, and 50-60%, respectively, as measured by recording Sr:GFP in 26 hAPF pupa. To evaluate the effect on tissue flow speed under such conditions we imaged the tissue, using autofocus detection (Metamorph software), at 15 min intervals (28-30 z-stack at 0.5 µm) typically for 20-22 h. To evaluate the efficacy of the *Shine-GAL4>sr*^*RNAi*^, after each experiment an image of Sr:GFP was acquired.

### Laser ablations

For large-scale ablations and analyses of tissue flow, we used an inverted spinning-disk confocal microscope (Roper/Zeiss) coupled to a UV (355 nm) ablation system. All other ablations were performed using a two-photon laser-scanning microscope (LSM 880 NLO, Zeiss): imaging was performed using mono-photon excitation, while local laser ablation was carried out using a biphoton Ti:Sapphire laser (MaiTai DeepSee, Spectra Physics) at 890 nm or 810 nm with <100 fs pulses and an 80 MHz repetition rate.

#### Large-scale repeated ablations

Pupae expressing Ecad:3xGFP or Ecad:GFP were imaged using a 40x/1.4 OIL DIC H/N2 PL FLUOR objective, with a time interval of 15 min and 60 z-steps of 1 µm each. Using the Ilas2 add-on in Metamorph software, a rectangular region was defined before 18 hAPF either over the neck region or within a region whose most anterior edge was aligned with the aDC macrochaetes (*x* = 0.4). Laser ablation was performed across 60 planes (with a z-step of 1 µm) every 3 hours starting at 18 hAPF using the UV 355 nm laser. The size of the ablation region and the repetition frequency were selected to prevent wound closure.

#### Ablation and protrusion stretching

Pupae expressing Ecad:3xGFP or Ecad:3xmKate2 (AJ signal) along with a mosaic labeling of LifeAct:Ruby or Myr:GFP (protrusion signal) at 24 hAPF were imaged using a 63x/1.4 OIL DICII PL APO objective. Imaging was restricted to a 100×100 pixel window (with a pixel size of 0.13 µm) using a bidirectional scan lasting 500 ms, and a z-stack of 4 planes (with a z-step of 0.5 µm), from the AJ level to the apical protrusion domain. Laser ablations, spanning approximately 2 cell diameters, along the ML axis and carried out with the biphoton Ti:Sapphire laser at 890 nm were performed at either the front or the back of LifeAct:Ruby-expressing clones, or at the front of Myr:GFP-expressing clones. Ablations were performed at a z-plane located 0.5 µm below the AJ plane, ensuring that the ablation region did not overlap with apical protrusions. Upon acquisition, the fluorescent signals were projected and segmented as described above (only planes situated above the AJ level were projected for protrusion signals). Junctions of interest were then manually selected and tracked at two timepoints: *t*_*initial*_ (immediately before ablation) and *t*_*final*_ (10 s after ablation). The area separating the junctions of interest between *t*_*initial*_ and *t*_*final*_ was measured (*Area*_*recoil*_). The spreading areas of the protrusion signal beyond the AJ signal were measured at *t*_*initial*_ (*Spreading*_*initial*_) and *t*_*final*_ (*Spreading*_*final*_). The percentage of stretching after ablation was calculated as 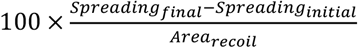

#### Muscle ablation

In the notum epithelium, the Dpy protein, produced by Sr-positive tendon cells, mediates adhesion between the tissue and the aECM^66,77,78^. In response to the pulling forces exerted by the dorsal-longitudinal indirect flight muscles (IFMs) from 26 hAPF onward, Dpy is required to maintain the bound between the tendon cells and the aECM, thereby preventing out-of-plane tissue deformations^58,66,79^. To avoid these out-of-plane deformations from confounding our measurements of tissue velocity, we first ablated the dorsal-longitudinal IFMs in control conditions and found that this ablation did not affect tissue velocity (Extended Data Fig. 2h). We therefore analyzed flow velocity in control and *Dpy*^*RNAi*^ conditions upon muscle ablation.

To ablate the dorsal-longitudinal IFM, the *Mhc-tau:GFP* line was used as a marker to visualize their precursors before 18 hAPF ^124^. Muscle ablation was performed prior to time-lapse imaging using a 40x/1.3 OIL DICII PL APO (UV) VIS-IR objective and the biphoton Ti:Sapphire laser. To ensure efficient muscle removal, the outline of each muscle precursor was manually traced to guide laser ablation specifically within the muscle, and ablation was repeated across multiple successive *z*-planes (typically 10) to ablate the entire muscle.

### Quantifications of Sr:GFP and Dpy:YFP level and domain size

Pupae expressing Ecad:3xmKate2 and either Sr:GFP or Dpy:YFP were imaged with 50 planes spaced by 1 µm *z*-step and 15 min time resolution typically between 17 and 30 hAPF. Upon defining the tissue *z*-map using the Ecad:3xmKate2 signal (see above), the Sr:GFP signal was measured from 5 to 15 µm below the AJ and the Dpy:YFP one was measured above the AJ. After rescaling and averaging the two hemi-thoraces, the Sr:GFP and Dpy:YFP fluorescence signals were temporally smoothed using a 1.25 h window. The Sr:GFP or Dpy:YFP signals were found to be minimal and not vary in *z* for the region between *x* = 0.4 and 0.55 this region was therefore used to measure background and its intensity subtracted for each pupae measurement. The resulting signal was then averaged over time between 24 and 28 hAPF and along the ML axis to measure their amplitude. For plotting the AP intensity profiles, the values were normalized by the mean peak value of the control. From these measurements the domain was located using the maximum intensity value and the edges of the domain were defined by the point at which the intensity was reduced to 20% of the maximal intensity. The distance between these points was used to measure the domain size.

### Statistics and reproducibility

Sample sizes vary in each experiment and animal samples were randomly selected within a given genotype for subsequent analyses. Experiments were repeated at least twice. Only animals correctly mounted for microscopy and without developmental delay were included in the analyses. The numbers of analyzed animals, ablations and cells are indicated in the figure legends or panels. For all graphs, error bars or shaded regions represent the standard deviation between animals, unless otherwise stated. In particular, for the stretching upon ablation and protrusion polarity the standard error of the mean between clones is shown. For all figures, representative microscopy images of at least two different experiments are shown. The statistical tests used to assess significance are stated in the figure legends and are two-sided. Statistical analyses were performed using SciPy^125^.

### Data, code, and materials availability

All datasets and reagents generated in this manuscript are available from the corresponding authors upon request. Scripts and codes used in this study are provided as a zip file and will be accessible at https://github.com/BellaicheLab/Notum-Flow-Scaling.

## Acknowledgments

We thank P. Adler, N. Dye, Y. Huang, E. Wieschaus, the Bloomington Drosophila Stock Center, Kyoto Stock Center, Transgenic RNAi Project at Harvard Medical School, and Vienna Drosophila Resource Center reagents; The Cell and Tissue Imaging Platform - PICT@BDD, member of the National Infrastructure France-BioImaging supported by the French National Research Agency (ANR-24-INBS-0005 FBI BIOGEN), for assistance with light microscopy; Mélina Durande for initial exploration in the theoretical modelling; Florencia di Pietro, Baptiste Tesson and Mari Yoshida for valuable comments on the manuscript. This work was supported by Institut Curie, CNRS and INSERM. ERC Advanced Scaling-Sensitivity (101020243); ARC (SL220130607097); ANR (TiMecaDiv 20CE13000801); ANR (Migrafold ANR-18-CE13-0021); ANR (ChronoDamage 20CE13-0013); CANCERO-INCA (PLBIO2020/BELLAICHE); ANR Labex DEEP (11-LBX-0044, ANR-10-IDEX-0001-02).

## Author contributions

Conceptualization: BG, AV, JD, FB, YB

Methodology: BG, AV, JD, FGr, FB, YB

Software: BG, AV, JD

Formal analysis: BG, AV, JD, FGä

Investigation: BG, AV, JD, FGä, LA, FB

Resources: IG

Data curation: BG, JD

Writing – original draft: BG, AV, JD, YB

Writing – review & editing: BG, AV, JD, FGä, LA, FGr, FB, YB

Visualization: BG, AV, JD, FB

Supervision: FB, YB

Project administration, Funding acquisition: YB

## Additional information

Supplementary Information is available for this paper.

## Extended Data Figures

**Extended Data Fig. 1.**
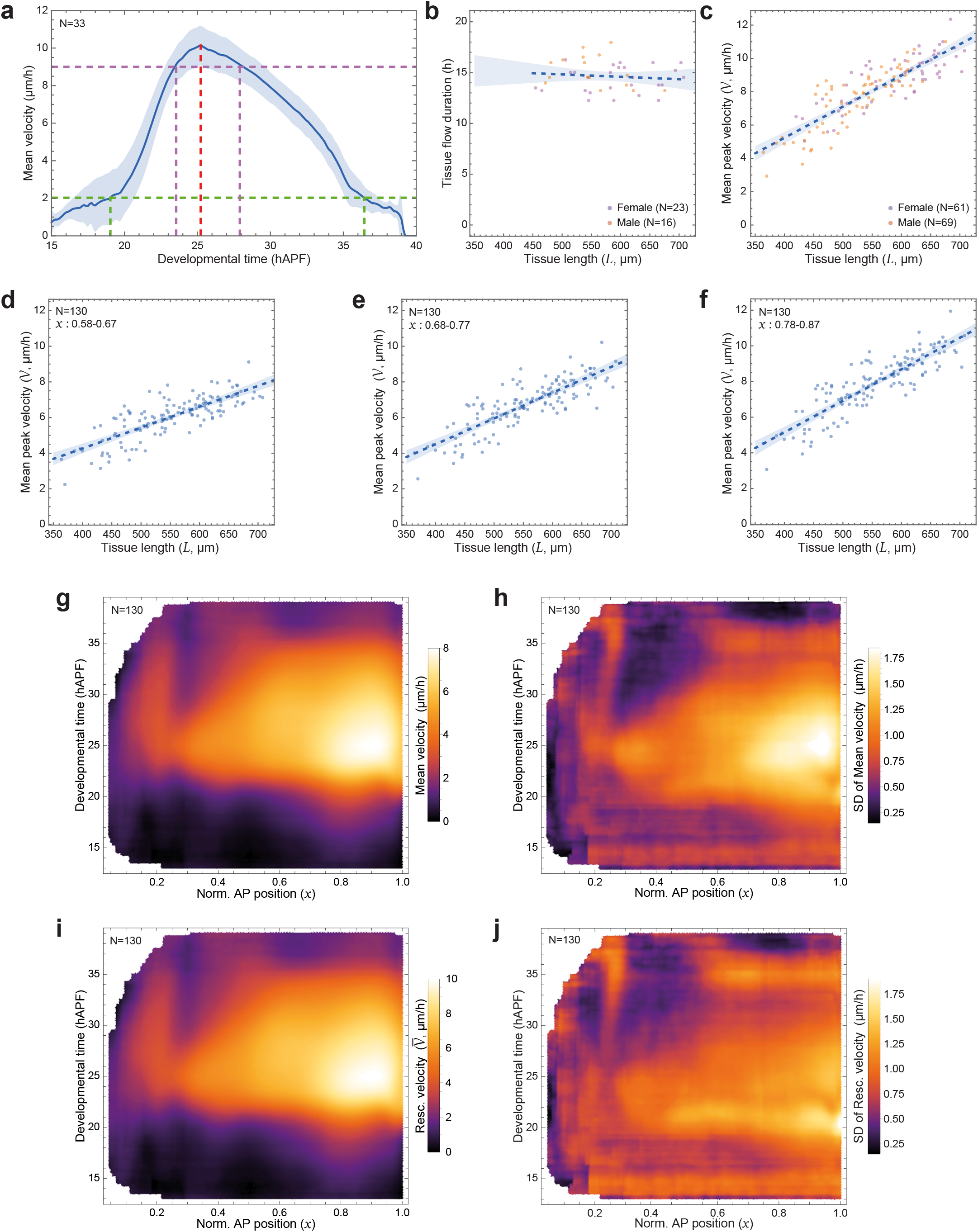
Thorax flow and the scaling of its speed with animal length. Unless otherwise stated: solid line, mean experimental data; dark blue dashed line, linear fit; shaded region, SD. N, number of animals. Genotypes are listed in Extended Data Table 3. 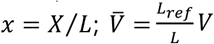. **(a)** Graph of the mean velocity (averaged from *x* = 0.88 to 0.97) as a function of the developmental time for commensurate tissue lengths (616 < *L* <707 µm). Green dashed lines, tissue flow onset (19 hAPF) and end (36 hAPF) as defined by 20% of the *V*_max_ (2.1 µm/h). Magenta dashed lines, peak of velocity onset (23.6 hAPF) and end (27.9 hAPF) as defined by 90% of the *V*_max_ (9.3 µm/h). Red dashed line, *V*_max_ = 10.34 µm/h (at 25.2 hAPF). Same data as Fig. 1c. **(b)** Graph of the tissue flow duration as a function of tissue length (*L*) for female and male pupae. Shaded area, 95% CI of the mean. R^2^= 0.012, *p*= 0.478 (Student’s *t*-test). Same fit and data as Fig. 1g. **(c)** Graph of the mean peak velocity (*V*) (velocity averaged from 24 to 28 hAPF and over 0.88 < *x* <0.97) as a function of tissue length (*L*) for female and male pupae. Shaded area, 95% CI of the mean. R^2^= 0.72, *p*= 2.94 × 10^−37^ (Student’s *t*-test). Same fit and data as Fig. 1h. **(d-f)** Graphs of the mean peak velocities (*V*) (velocity averaged from 24 to 28 hAPF) as a function of tissue lengths (*L*) at various AP (*x*) positions; averaged over 0.58< *x* <0.67 (d), 0.68< *x* <0.77 (e), and 0.78< *x* <0.87 (f). Shaded area, 95% CI of the mean. R^2^= 0.74 (d); 0.68 (e); 0.62 (f). *p*= 2.2 × 10^−39^ (d); 1.9 × 10^−33^ (e); 8.9 × 10^−29^ (f) (Student’s *t*-test). **(g-h)** Kymograph of the mean velocity (averaged across tissues) (g) and SD (h) along the normalized AP position (*x*) as a function of developmental time for all tissue lengths (*L*). Same data as Fig. 1h. **(i-j)** Kymograph of the mean rescaled AP velocity (averaged across tissues) (i) and SD (j) as a function of the normalized AP position (*x*) and developmental time for all tissue lengths (*L*). Same data as Fig. 1h.

**Extended Data Fig. 2.**
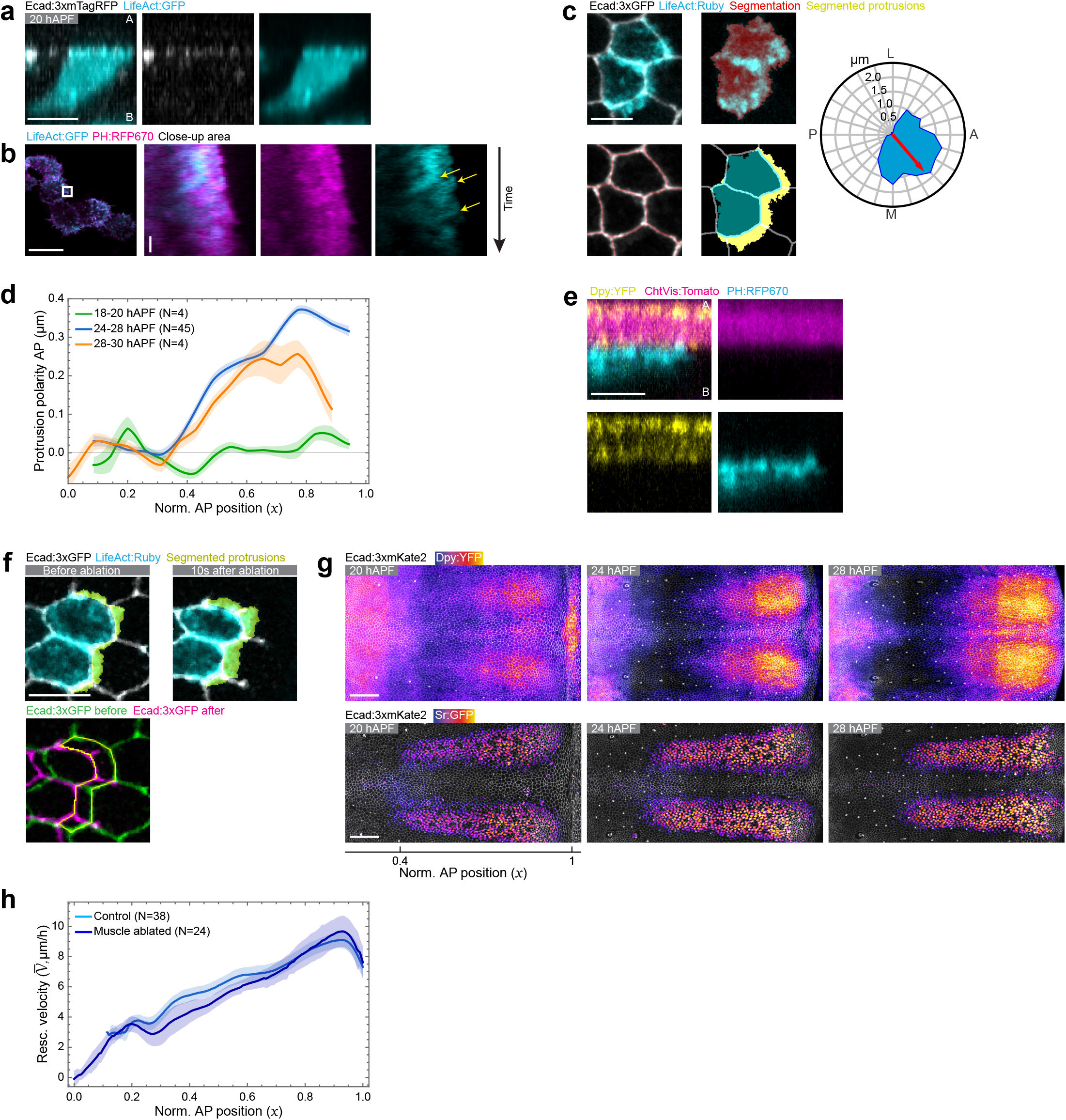
Regulation of migration flow during notum development. The AP axis is horizontal with the posterior to the left. When relevant, the normalized AP position (*x*) is indicated below the images. Quantifications are averaged from 24 to 28 hAPF. Unless otherwise stated: Images were acquired in the anterior notum region at 25 hAPF; solid line, mean experimental data; shaded region, SD. N, number of animals. Genotypes are listed in Extended Data Table 3. 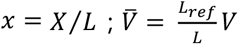. **(a)** Apical-basal section along the AP axis of a live notum expressing Ecad:3xTagRFP at 20 hAPF upon LifeAct:GFP clonal expression. A, Apical; B, Basal. **(b)** LifeAct:GFP, PH:RFP670 notum live images upon LifeAct:GFP and PH:RFP670 clonal expression and corresponding kymographs from Supplementary Video 5. The kymograph was generated by projecting the signal within the white square perpendicularly to the protrusion direction. Yellow arrows, LifeAct:GFP retrograde flow within the protrusion. Same clone as Fig. 2c. **(c)** Ecad:3xGFP notum live images at 24 hAPF upon LifeAct:Ruby clonal expression (top left). Segmentation of the LifeAct:Ruby and associated fluorescent signal (top right). Segmentation of the Ecad:3xGFP and associated fluorescent signal (bottom left). Overlay of both segmentations and resulting segmented protrusions defined as the part of the LifeAct:Ruby segmented signal outside the segmented AJ signal (bottom right). Rose plot of the clone polarity signal (blue area) and the resulting polarity vector (red arrow). The orientation of the polarity vector is the main direction of protrusion spreading, while its length corresponds to the amplitude of polarization. **(d)** Graph of the protrusion polarity AP component (mean ± SEM) as a function of the normalized AP position (*x*) averaged from 18 to 24, 24 to 28, and 28 to 30 hAPF. **(e)** Apical-basal section along the AP axis of a live notum expressing Dpy:YFP (bottom left), ChtVis:Tomato (top right), and clonally PH:RFP670 (bottom right), with the merged shown in the top left. A, Apical; B, Basal. **(f)** Ecad:3xGFP live notum image at 24 hAPF upon LifeAct:Ruby clonal expression with segmented protrusions before (top left) and 10 s after (top right) anterior AJ ablation. Overlay of both Ecad:3xGFP images and the AJ recoil induced by the ablation (yellow outline) used for normalization (see Methods). **(g)** Ecad:3xmKate2 notum live images showing Dpy:YFP (top) and Sr:GFP (bottom) intensities at 20, 24, and 28 hAPF. Dpy:YFP and Sr:GFP are projected above the AJ and at the level of the nucleus, respectively. **(h)** Graph of the mean rescaled velocity 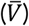 as a function of the normalized AP position (*x*) in control tissue, and tissue after dorsal-longitudinal indirect flight muscles (IFM) ablation (see Methods). From 26 hAPF onwards, IFM attach to the tendon cells, exerting pulling forces on the epithelial cells and leading to epithelial folding in the most anterior part of the notum in absence of Dpy. Ablation of the dorsal-longitudinal IFMs does not affect tissue flow velocity and was therefore used in Fig. 2h to measure tissue flow velocity in absence of folding. Scale bars: 100 µm (g), 10 µm (a, b, c, e, f), 30 s (Kymograph b).

**Extended Data Fig. 3.**
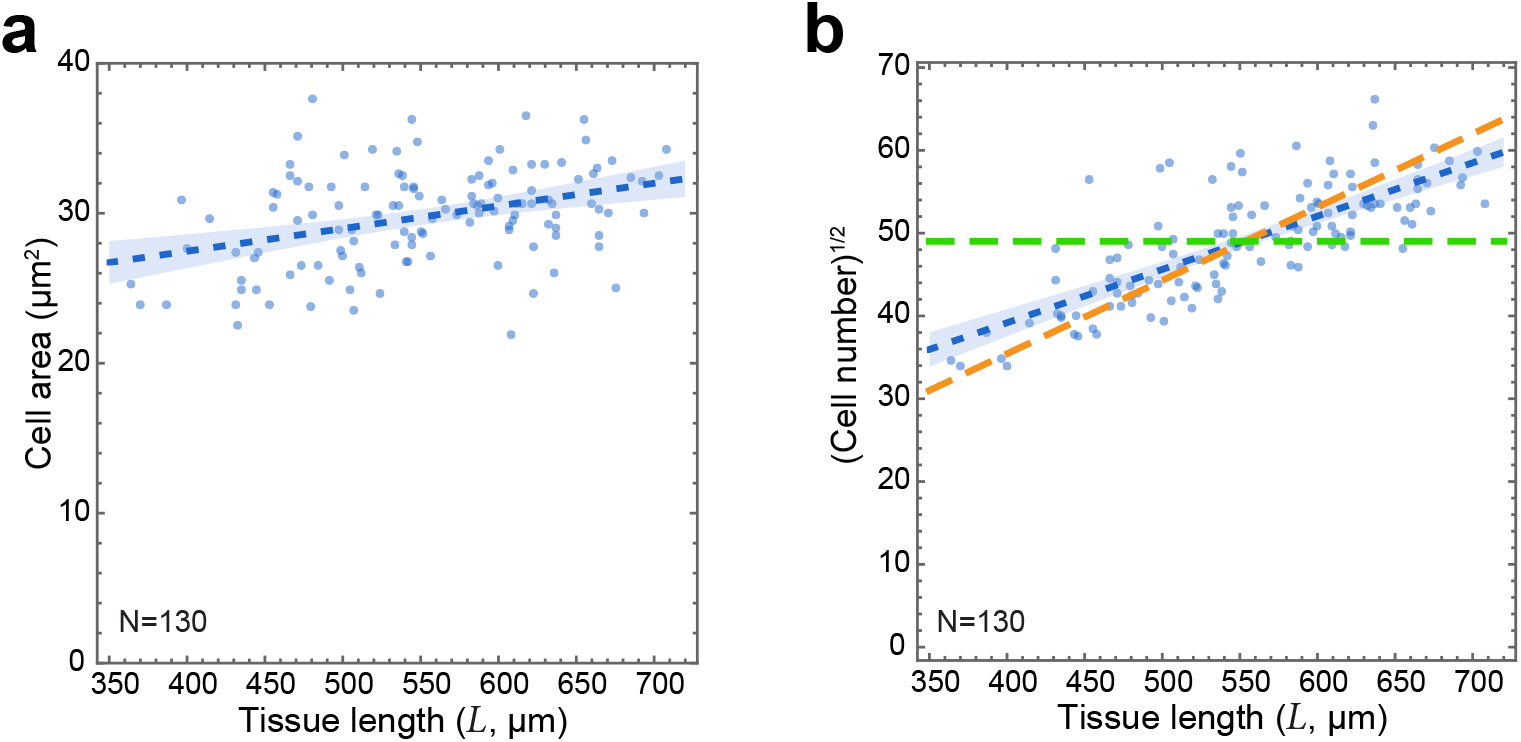
Variation in cell number and cell apical area as a function of tissue length. Quantifications were performed at 25 hAPF. *p* value, Student’s *t*-test. Shaded area, 95% CI of the mean. Unless otherwise stated: Dark blue dashed line, linear fit. N, number of animals. Genotypes are listed in Extended Data Table 3. **(a)** Graph of the mean cell area as a function of tissue length (*L*). Mean cell areas were determined by segmenting a strip of tissue located between *x* = 0.43 and *x* = 0.89 along the AP axis, and between 50% and 70% of the width of the ML axis (between the midline and one aDC macrochaetae, see Fig. 1b) in each animal. R^2^= 0.14, *p*= 1.2 × 10^−5^. Same data as Fig. 1h. **(b)** Graph of the square root of cell number as a function of tissue length (*L*). The cell number was estimated in a strip centered on the midline with the same AP length as (a), but spanning the whole ML axis. This ROI area was divided by the mean cell area to estimate the cell number. Green dashed line, cell number if the ROI size was purely accompanied by a change in cell area; orange dashed line, cell number if the ROI size was purely accompanied by a change in cell number. R^2^= 0.59, *p*= 6.1 × 10^−26^. Same data as Fig. 1h.

**Extended Data Fig. 4.**
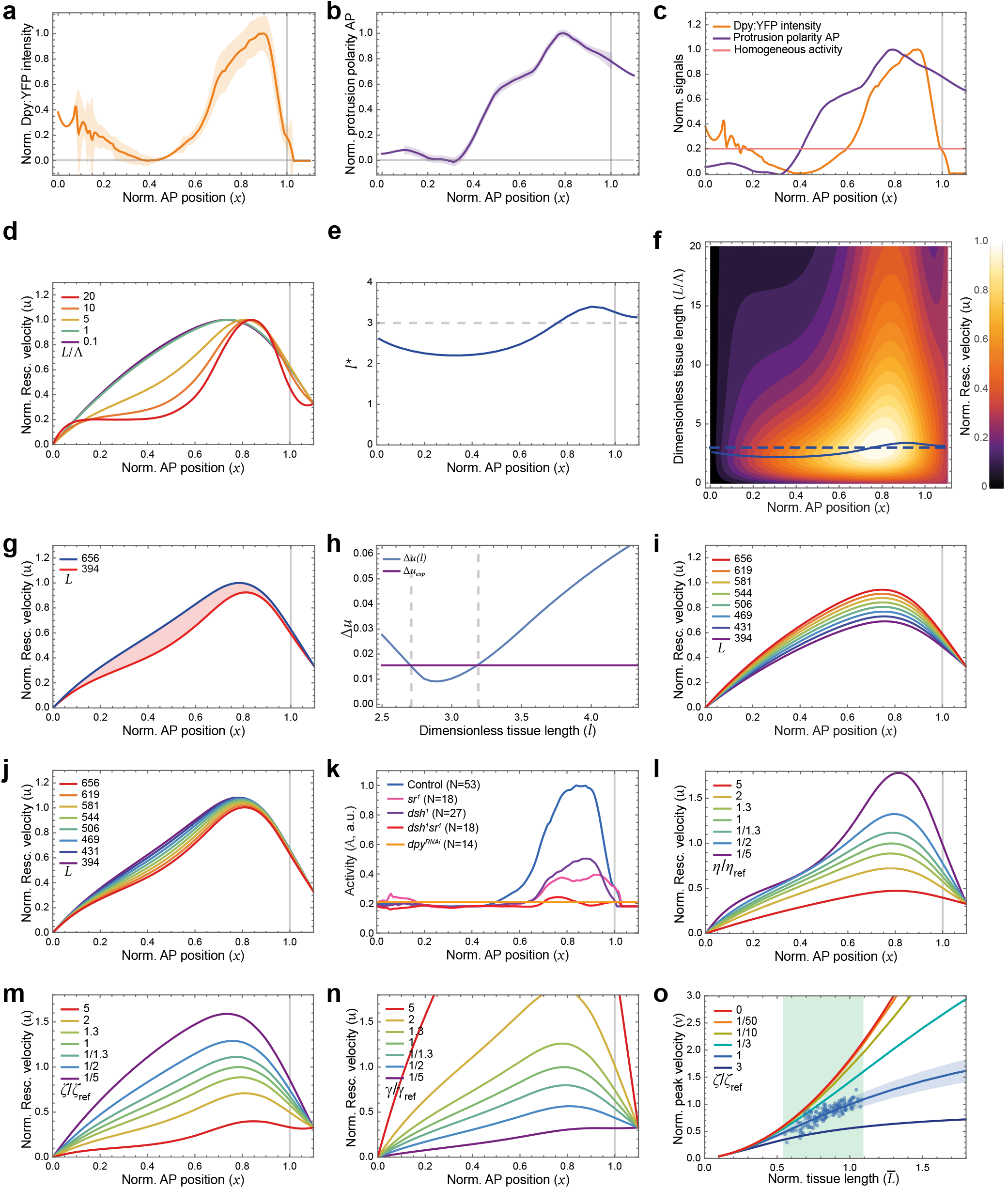
Modelling of collective migration. The AP axis is horizontal with the posterior to the left. Quantifications are averaged from 24 to 28 hAPF. Unless otherwise stated: solid line, mean experimental data; shaded region, SD; vertical gray line (*x* >1), anterior limit of data (beyond this point data is extrapolated). N, number of animals. 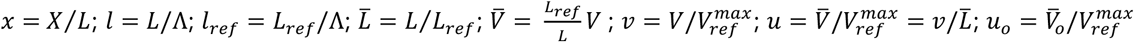. **(a)** Graph of the mean normalized Dpy:YFP intensity as a function of the normalized AP position (*x*). Same data as Fig. 2m. **(b)** Graph of the normalized protrusion polarity AP component (mean ± SEM) as a function of the normalized AP position (*x*). Same data as Fig. 2n. **(c)** Graph overlay of the normalized Dpy:YFP intensity (same as a), normalized protrusion polarity (same as b) and homogeneous activity (*Ao*) as a function of the normalized AP position (*x*). SD are not displayed for clarity. **(d)** Graph of the model predictions for the normalized rescaled velocity (*u*) as a function of the normalized AP position (*x*) for different values of *l*. The parameter *u*_2_ was adjusted so that *u*_*max*_ = 1. **(e)** Graph of the dimensionless tissue length (*l*) value at which the normalized rescaled velocity (*u*) admits a maximum (*l*^∗^) as a function of the normalized AP position (*x*) with *u*_2_ =10. Horizontal grey dashed line, *l* = *l*_*ref*_ = 3. **(f)** Heatmap of the normalized rescaled velocity (*u*) as a function of the normalized AP position (*x*) and of the dimensionless tissue length (*l*) with *u*_*o*_ =10. Blue solid line, *l*^∗^(*x*); Blue dashed line, *l* = *l*_*ref*_ = 3. Curves are replotted from E. **(g)** Graph of the model prediction of the normalized rescaled velocity (*u*) as a function of the normalized AP position (*x*) for the smallest and largest tissue length bins (for *l*_*ref*_ = 5). Light red, area between the curves. **(h)** Graph of the model prediction of (Δ*u*(*l*)) as a function of the dimensionless tissue length (*l*). Δ*u*_*exp*_ (see theory note § 2.9) reaches a minimum in the vicinity of *l* = 2.9 ≃ 3, corresponding to the best overlap of the theoretical curves in the interval 0 < *x* < 0.7 in Fig. 4g. (**i-j**) Graph of the model prediction for the normalized rescaled velocity (*u*) as a function of the normalized AP position (*x*) with *u*_*o*_ = 8.6 and *Λ* values set such that *l*_*ref*_ =1 (i) or *l*_*ref*_ = 5 (j). (**k**) Graph of the activity (*A*) as a function of the normalized AP position (*x*) in control and mutant tissues used in the model. Each *A*(*x*) function was built for each condition based on their AP profiles of Dpy:YFP intensity (see Fig. 2m) and protrusion polarity (see Fig. 2i,n and Extended Data Fig. 4a-c). Estimated SD are not displayed for clarity. (**l-n**) Graphs of the model prediction for the normalized rescaled velocity (*u*) as a function of the normalized AP position (*x*) with evolving parameters *η* (l), *ζ* (m) and *γ* (n). The indicated numbers and fractions correspond to the ratio of the parameter over its control reference value. In all cases *u*_*neck*_ was kept constant. (**o**) Graph of the model prediction for the normalized velocity (*v*) as a function of the normalized tissue length 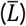 for different values of *ζ*/*ζ*_*ref*_. Light green vertical strip, range of physiological tissue lengths, including the linear domain of the *ζ*/*ζ*_*ref*_ = 1 curve (located in the vicinity of 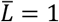). Lowering *ζ*/*ζ*_*ref*_ shifts the curves towards higher 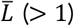, away from the physiological tissue length range (linearity of the scaling 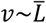 is gradually lost and gets closer to 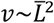. Increasing *ζ*/*ζ*_*ref*_ shifts the linear domain of the curves towards lower 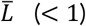, away from the physiological tissue length range (linearity of the scaling is gradually lost and gets closer to *v~cst)*. Same data as Fig. 1h; *ζ*/*ζ*_*ref*_ = 1 curve is replotted from Fig. 4i.

**Extended Data Fig. 5.**
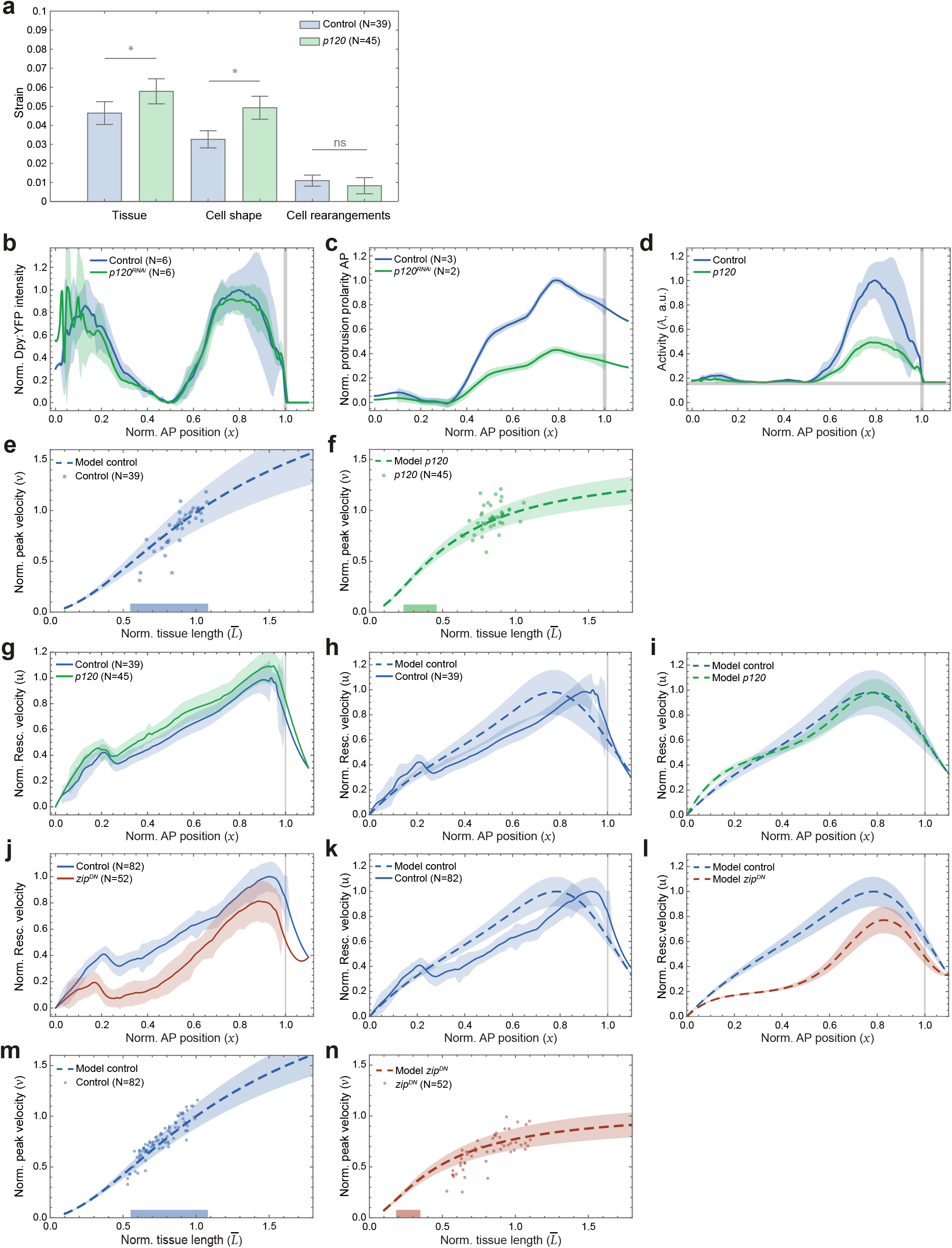
Characterization of tissue dynamics in *p120* and *zip*^*DN*^. The AP axis is horizontal with the posterior to the left. Quantifications are averaged from 24 to 28 hAPF. Unless otherwise stated: solid line, mean experimental data; dashed line, model; shaded region, SD. N, number of animals. Genotypes are listed in Extended Data Table 3. 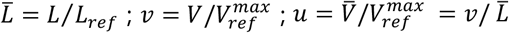. **(a)** Plot of the cumulated tissue strain (traceless part representing pure contraction-elongation along the AP axis) and corresponding strains for cell shape changes and cell rearrangements in control and *p120* tissues. Bars represent the cumulated pure shear (deviatoric part of strain tensor) quantifying tissue elongation (with no change in area) along the AP axis, and the respective contributions of cell shape changes and cell rearrangements. Tissue strain *p*= 0.0398, Cell shape *p*= 0.0175, Cell rearrangements *p*= 0.0681 (Mann-Whitney U); \**p*< 0.05; ns, not significant. **(b)** Graph of the mean normalized Dpy:YFP intensity as a function of the normalized AP position (*x*) at 25 hAPF in control and *p120*^*RNAi*^ tissues. **(c)** Graph of the normalized protrusion polarity AP component (mean ± SEM) as a function of the normalized AP position (*x*) in control and *p120*^*RNAi*^ tissues. **(d)** Graph of the activity (*A*) (based on b and c) as a function of the normalized AP position (*x*) in control and *p120*^*RNAi*^ tissues. (**e**,**f**) Graph of the model prediction for the normalized velocity (*v*) as a function of the normalized tissue length 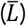 for control (e) and *p120* (f) tissues. Data point velocities and tissue lengths were rescaled by the mean velocity in tissues of commensurate lengths (*V* = 10.6 μm/h for animal length range: 623 µm < *L* < 704 µm) and the reference animal size *L*_*ref*_ = 656 μm. Horizontal bars on the abscissa, position of the linear domain of the model extended to the physiological length range in control (e) and its corresponding location and width in the model *p120* curve (f). Same data as Fig. 5a. **(g)** Graph of the normalized rescaled velocity (*u*) as a function of the normalized AP position (*x*) in control and *p120* tissues of commensurate lengths (622 µm < *L* < 704 µm). Same data as Fig. 5b. **(h)** Graph of the normalized rescaled velocity (*u*) as a function of the normalized AP position (*x*) in control tissues of commensurate lengths (623 µm < *L* < 704 µm) and the associated model prediction. Same data as Fig. 5b. **(i)** Graph of the model predictions for the normalized rescaled velocity (*u*) as a function of the normalized AP position (*x*) in control and *p120* tissues. **(j)** Graph of the normalized rescaled velocity (*u*) as a function of the normalized AP position (*x*) in control and *zip*^*DN*^ tissues of commensurate lengths (615 µm < *L* < 713 µm). Same data as Fig. 5e. **(k)** Graph of the normalized rescaled velocity (*u*) as a function of the normalized AP position (*x*) in control tissues of commensurate lengths (615 µm < *L* < 663 µm) and the associated model prediction. Same data as Fig. 5e. **(l)** Graph of the model predictions for the normalized rescaled velocity (*u*) as a function of the normalized AP position (*x*) in control and *zip*^*DN*^ tissues. (**m, n**) Graph of the model prediction for the normalized velocity (*v*) as a function of normalized tissue length 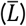 for control (m) and *zip*^*DN*^ (n) tissues. Data point velocities and tissue lengths were rescaled by the mean velocity in tissues of commensurate lengths (*V* = 9.9 μm/h for animal length range: 615 µm < *L* < 663 µm) and the reference animal size *L*_*ref*_ = 656 μm. Horizontal bars on the abscissa, position of the linear domain of the model extended to the physiological length range in control (m) and its corresponding location and width in the model *zip*^*DN*^ curve (n). Same data as Fig. 5d.

## Extended Data Tables

**Extended Data Table 1.** Variables used for quantification and 1D modeling of collective cell migration.

| symbol | name/description | definition | approx. largest range/value | units |
| --- | --- | --- | --- | --- |
| $L$ | notum length | | [350, 730] | $\mu\text{m}$ |
| $X$ | position on AP axis | | [0, L] | $\mu\text{m}$ |
| $V$ | velocity | | [0, 10] | $\mu\text{m}/\text{h}$ |
| $L_{ref}$ | Mean $L$ for reference animals | | 656 | $\mu\text{m}$ |
| $V_{ref}$ | Mean $V$ for reference animals | | [0, 10] | $\mu\text{m}/\text{h}$ |
| $V_{ref}^{max}$ | $V_{ref}$ max value | | 9.9 | $\mu\text{m}/\text{h}$ |
| $x$ | rescaled AP axis position | $X/L$ | [0, 1] | |
| $v$ | normalized velocity | $V/V_{ref}^{max}$ | [0, 1] | |
| $\bar{V}$ | rescaled velocity | $\frac{L_{ref}}{L}V$ | | $\mu\text{m}/\text{h}$ |
| $u$ | normalized rescaled velocity | $\bar{V}/V_{ref}^{max}$ | [0, 1] | $\text{h}^{-1}$ |
| $\eta$ | tissue viscosity | | | $\text{N m}^{-1} \text{h}$ |
| $\zeta$ | friction on ECMs | | | $\text{N m}^{-3} \text{h}$ |
| $\Lambda$ | characteristic hydrodynamic length | $(\eta/\zeta)^{1/2}$ | 220 | $\mu\text{m}$ |
| $\bar{V}_o$ | characteristic rescaled velocity | $\frac{\gamma L_{ref}}{(\eta\zeta)^{1/2}}$ | | $\mu\text{m}/\text{h}$ |
| $u_o$ | characteristic normalized rescaled velocity | $\bar{V}_o/V_{ref}^{max}$ | | $\text{h}^{-1}$ |
| $l$ | dimensionless notum tissue length | $L/\Lambda$ | | |
| $\bar{L}$ | normalized notum tissue length | $L/L_{ref}$ | [0.5, 1.1] | |
| $l^*$ | $l$ value at which $u(x, l)$ reaches a maximum | $\partial_l u(x, l)\vert _{l^*} = 0$ | | |

**Extended Data Table 2.** *Drosophila* stocks used in this study.

| *Drosophila* stocks | References | Availability |
| --- | --- | --- |
| *Ecad:GFP* | 126 | Gift from Y. Huang |
| *Ecad:3xGFP* | 113 | Upon request |
| *Ecad:3xmKate2* | 113 | Upon request |
| *Ecad:3xmTagRFP* | 113 | Upon request |
| *Dpy:YFP* | 127 | Kyoto Stock Center #115238 |
| *Sr:GFP* | This study | Upon request |
| *Mhc-tau:GFP* | N/A | Bloomington Stock Center #38460 |
| *Act5C-GAL4* | N/A | Bloomington Stock Center #3954 |
| *Shine-GAL4* | 85 | Bloomington Stock Center #94199 |
| *Act-FRT-y<sup>+</sup>-FRT-GAL4* (Chr II, III) | N/A | Bloomington Stock Center #3953 and #4413 |
| *Act-FRT-CD2-FRT-GAL4* | 128 | Bloomington Stock Center #4780 |
| *Tub-GAL80<sup>ts</sup>* | 129 | Bloomington Stock Center #7017 |
| *UAS-LifeAct:Ruby* | N/A | Bloomington Stock Center #35545 |
| *UAS-LifeAct:GFP* | N/A | Bloomington Stock Center #35544 |
| *UAS-ChtVis:Tomato* | 68 | Gift from P. Adler |
| *UAS-PH:RFP670* | This study | Upon request |
| *UAS-FRT-STOP-FRT-Myr:GFP* | N/A | Bloomington Stock Center #55811 |
| *UAS-CAAX:TagBFP* | 130 | Bloomington Stock Center #55090 |
| *UAS-dpy<sup>RNAi</sup>* | 131 | Vienna Drosophila Resource Center #44029 |
| *UAS-LacZ* | N/A | Bloomington Stock Center #8529 |
| *UAS-w<sup>RNAi</sup>* | 132 | Bloomington Stock Center #33644 |
| *UAS-sr<sup>RNAi</sup>* | 131 | Vienna Drosophila Resource Center #105282 |
| *UAS-p120<sup>RNAi</sup>* | 132 | Bloomington Stock Center #51729 |
| *UAS-zip<sup>DN</sup>* | 133 | Gift from E. Wieschaus |
| *dsh<sup>1</sup>* | 134 | Bloomington Stock Center #5298 |
| *sr<sup>1</sup>* | 135 | Bloomington Stock Center #598 |
| *p120<sup>308</sup>* | 136 | Gift from N. Dye |
| *hs-FLP* (Chr I, II) | 137 | Bloomington Stock Center #8862 and #1813 |

**Extended Data Table 3.**
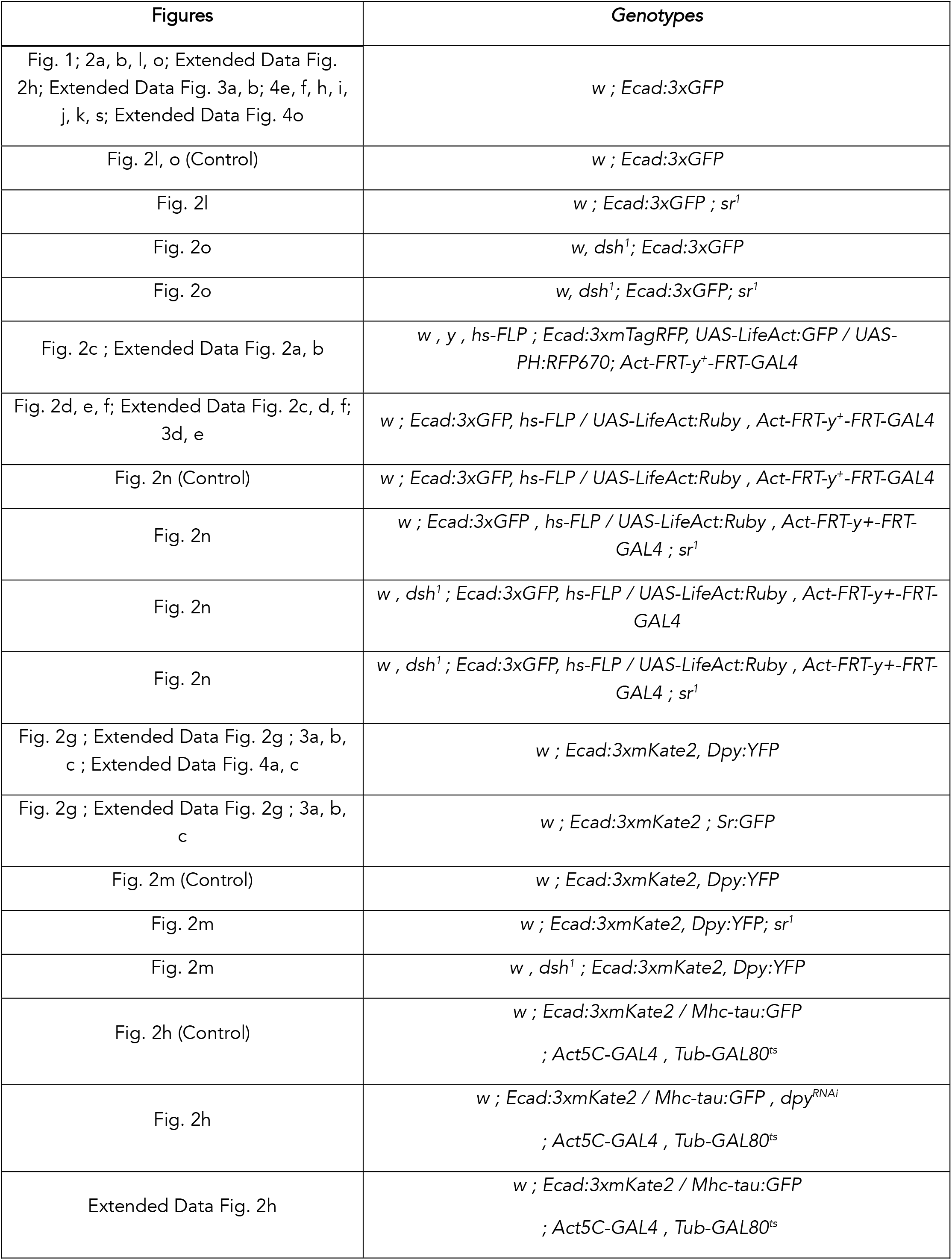
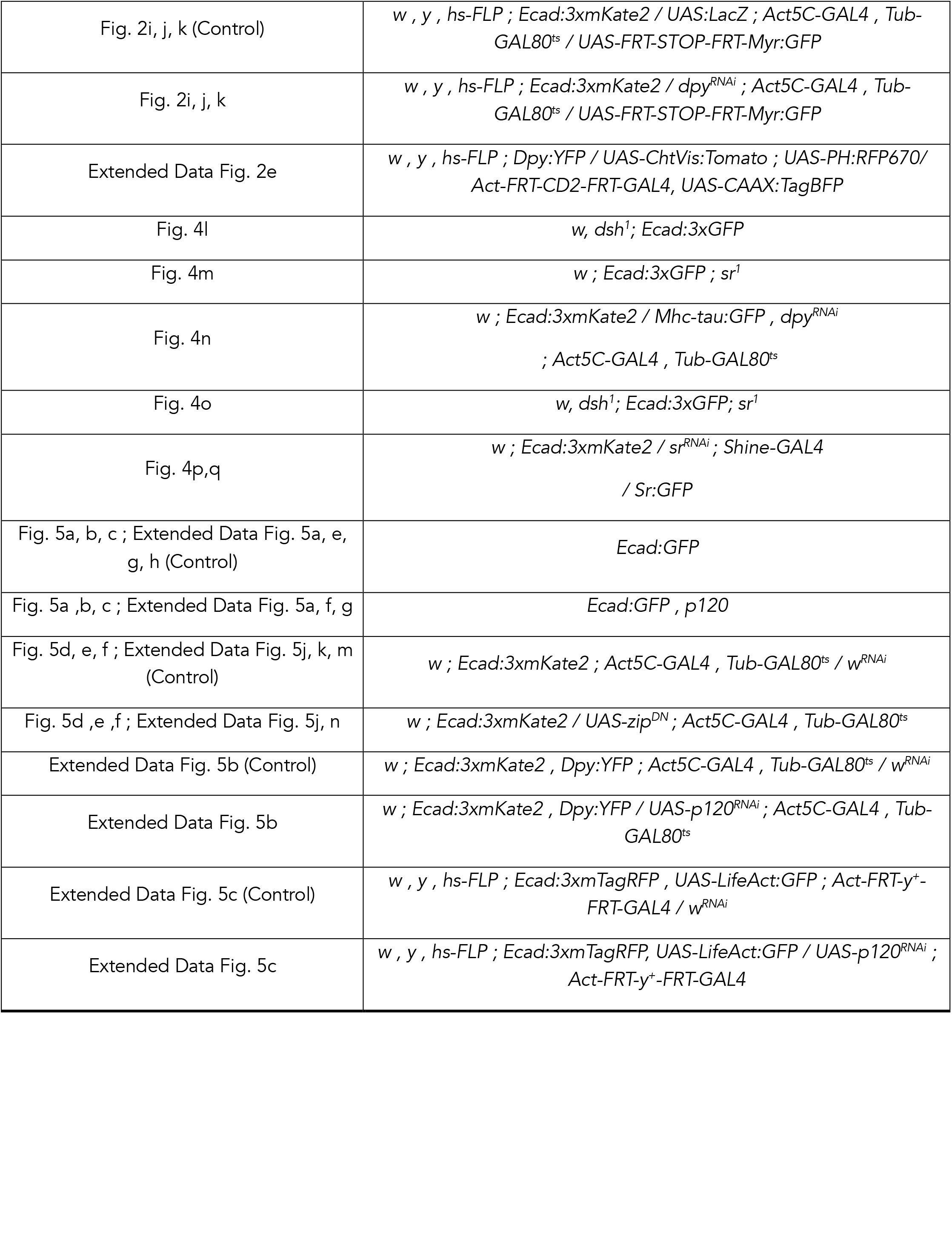
Experimental genotypes.

## Supplementary Information for

## Legends for Supplementary Videos

**Supplementary Video 1. Time-lapse movie of notum collective flow**.

Ecad:3xGFP time-lapse of the medial notum region with a tissue length of *L* = 673 µm between 16:30 and 31:30 hAPF. Frame rate: 1 frame / 15 min. Anterior to the right. Scale bar: 50 µm.

**Supplementary Video 2. Time-lapse movie of notum collective flow for different notum sizes**.

Side-by-side time-lapse images of Ecad:3xGFP in the notum for different tissue lengths (*L* = 345, 514 and 726 µm) between 22 and 28 hAPF. Frame rate: 1 frame / 15 min. Anterior to the right. Scale bar: 50 µm.

**Supplementary Video 3. Time-lapse movie of notum collective flow upon repeated laser ablation at the neck**.

Ecad:3xGFP time-lapse of the notum before and after repeated neck ablation between 15:15 and 28 hAPF. Orange dashed box: ablated region. Ablations were repeated every 3 hours. Frame rate: 1 frame / 15 min. Anterior to the right. Scale bar: 50 µm.

**Supplementary Video 4. Time-lapse movie of notum collective flow upon repeated laser ablation near the anterior DC macrochaetae**.

Ecad:3xGFP time-lapse of the notum before and after repeated posterior ablation (behind the anterior DC macrochaetae) between 15:45 and 28 hAPF. Orange dashed box: ablated region. Ablations were repeated every 3 hours. Frame rate: 1 frame / 15 min. Anterior to the right. Scale bar: 50 µm.

**Supplementary Video 5. Time-lapse movie of F-Actin and membrane in migrating notum cells**.

Time-lapse of the apical cell region in the anterior notum migration domain upon LifeAct:GFP and PH:RFP670 clonal expression and close-up (white box). Frame rate: 1 frame / 3 seconds. Anterior to the right. Scale bar: 5 µm.

## Theory Note

In this Theory Note, we detail the theoretical approach used to model collective cell migration along the Antero-Posterior (AP) axis dorsal thorax (notum) that occurs during the pupal development of *Drosophila*, and to study its observed scaling with tissue size.

We first provide general considerations on the scaling of tissue flows with tissue size (section 1), then we introduce a minimal model of cell migration that we solve, compare to experimental data, and use to make new predictions (section 2).

## 1 General Considerations

### 1.1 Definitions

A key quantity when studying animal development and cell displacement is the tissue flow velocity vector in the laboratory referential **V**(**X**) at a given position **X**. We take the origin of our axes at one end of our tissue, the rear of the notum in our case, and therefore **X** does not locate the same part of the tissue whether we consider a large animal or a small one. It is therefore natural to introduce the dimensionless position **x** = **X***/L*, where *L* is a length characterizing the tissue size (here simply the notum length) that locates specific parts of a tissue regardless of its size and that we can make appear in **V**:

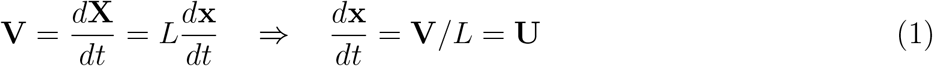

where **U** is the flow velocity rescaled by the notum length *L* that we refer to as “rescaled velocity” in the following. It represents the *relative* displacement of a piece of tissue during *dt* regardless of the tissue size. **U** is homogeneous to the inverse of a time and is the relevant quantity to study in what follows. Indeed, in addition to naturally appear when using **x**, it is also relevant to study the scaling of cell migration, since for a given developmental period, cell must travel the same *relative* distance in the tissue to end up at the correct location. This ideally means keeping the same rescaled velocity *U*, regardless of the tissue size *L*. Note that in the main text we also defined

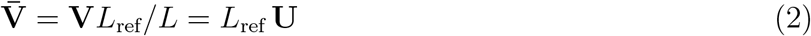

(with *L*_ref_ = 656 *µm* being the length of a tissue of reference size) which is therefore directly proportional to **U** but keeps the units of a velocity. Since we are only interested in the component of the velocity along the AP axis and its variation along the same axis, we orient our axes to make the *x* and AP axes coincide and we write everything in 1D for simplicity. We assume the dimensionless AP position *x* to vary between 0 and *x*_*neck*_ ≳ 1 and the tissue length *L* to vary within the range *L*_min_ and *L*_max_. For convenience, we also define the normalized tissue length 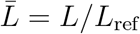.

### 1.2 Perfect Scaling of Tissue Flow at All Scales

The tissue flow velocity *V* being proportional to the tissue size *L* is equivalent to the rescaled velocity *U* being independent of the tissue size *L* (Eq. 1), which mathematically translates to:

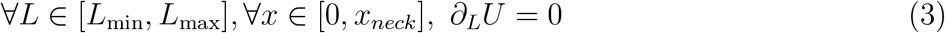

which is equivalent to:

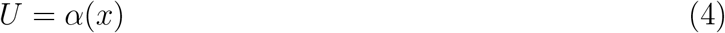

where *α* is a function solely depending on *x*. Coming back to the velocity, global scaling at all scales therefore occurs for:

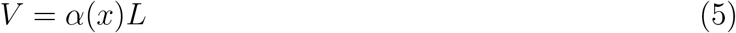

Thus, a global scaling of tissue flow at all sizes in [*L*_min_, *L*_max_] can occur if and only if *V* = *α*(*x*)*L*. If this size range spans several orders of magnitude, the scaling could then occur at different scales of tissue sizes. Note that in this case, the velocity *V* perfectly scales linearly with *L* at all *x* locations on the AP axis.

### 1.3 Scaling of Tissue Flow in a Limited Range of Sizes

A weaker requirement to achieve size-independence of rescaled tissue flow *U* would be if *U* was not globally independent of *L* on the full size range like before, but only locally independent, namely if *U* admitted an extremum at a given *L*^∗^, which translates to:

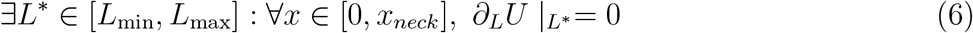

This means that in the vicinity of length *L*^∗^, the rescaled velocity *U* is independent of *L*. Therefore, for tissues whose lengths remain close to *L*^∗^, their flow will scale linearly with *L*.

The physiological range of tissue sizes (a subpart of [*L*_min_, *L*_max_]) should therefore contain *L*^∗^ and not extend too far around it for the tissue flow velocity to remain proportional to its size in good approximation. The quality of the linear scaling of tissue flow around *L*^∗^ will therefore depend on how *U* reaches its maximum at *L*^∗^ (namely how peaked it is) and how wide is the physiological range of sizes around it. If the *U*(*L*) curve remains rather flat around its maximum over this range, the linear scaling will occur in the whole physiological range in good approximation. In contrast, if *U*(*L*) is rather peaked around its maximum, the scaling will only occur in the near vicinity of *L*^∗^ and sizes near the physiological range boundaries will depart from linear scaling.

Note that in this case as well, this time only for tissue sizes in the vicinity of *L*^∗^, the velocity scales linearly with *L* at all *x* locations on the AP axis.

### 1.4 Approximate Scaling of Tissue Flow in a Limited Range of Sizes

An even weaker requirement would be that *L*^∗^ would weakly depend on *x*, namely that the length at which *U* reaches its extremum slightly varies along the x axis:

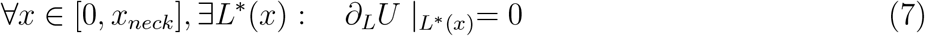

In that case, we would therefore still get a constant rescaled tissue flow *U* for tissue sizes in the vicinity of *L*^∗^(*x*), and equivalently, a tissue flow *V* linearly scaling with *L*, but *not* across the whole AP axis. Indeed, for a physiological range of sizes centered around *L*^∗^(*x*_*s*_), *x*_*s*_ being a particular location on the AP axis, tissue sizes will therefore display linear *V* (*L*) relationship across tissue sizes around *L*^∗^(*x*_*s*_) (similarly to section 1.3), but at other locations *x*_*o*_ ≠ *x*_*s*_, *V* (*L*) may depart from this linear relationship as *L*^∗^(*x*_*o*_) may be closer the boundaries of the physiological size range or even be out of it. In other words, the physiological size range cannot simultaneously be centered around *L*^∗^(*x*_*s*_) and *L*^∗^(*x*_*o*_) if those lengths are too different, and some part of the AP axis will therefore display better linear scaling than others. Nevertheless, if *L*^∗^ weakly depends on *x*, this case may end up landing quite close to section 1.3 (as *L*^∗^(*x*_*o*_) ≃ *L*^∗^(*x*_*s*_)).

### 1.5 Conclusion

In the three previous cases we saw that linear scaling of tissue flow *V* with tissue length *L* (or equivalently independence of *U* = *V/L* with respect to *L*) could either occur: (i) globally, namely at all tissue sizes, and even across different scales, and homogeneously along the AP axis (section 1.2); (ii) only locally and approximately, namely around a range of tissue sizes (centered on *L*^∗^), the smaller and the bigger tissues in the physiological size range departing from linearity, and homogeneously along the AP axis (1.3); (iii) locally and even more approximately, as in addition to departure from linear scaling for tissue sizes at the boundaries of the size range, the linear scaling also cannot perfectly occur along the whole AP axis, thereby resulting in some parts of the tissue displaying better linear scaling than others (section 1.4).

## 2 Minimal Model of Cell Migration

### 2.1 Introduction

With these considerations in mind, and by making common assumptions, we now aim at elaborating a simple physical model describing collective cell migration in the *Drosophila* notum that only incorporates the main ingredients, a classic approach in modeling^1–15^. Given the complexity of developing living tissues, our main goal is to obtain a qualitative agreement with experimental data and to understand the physical mechanism at the core of this scaling of velocity with animal size. On the one hand the tissue thickness (in the z direction) is much smaller than its two other dimensions at the apical surface (x, y) (Extended Data Fig. 2a), and on the other hand, the tissue active flow occurs very predominantly along the AP axis (x direction, Fig. 1b,f), we therefore have *V*_*x*_ ≫ *V*_*y*_, *V*_*z*_, and the two latter flows are negligible with respect to the flow along the AP axis direction.

Since we are only interested in describing the velocity component along the AP axis, corresponding to the *x* axis, namely *V*_*x*_ that we simply write *V* in what follows, and its spatial evolution along this same axis, we will derive here a 1D model for simplicity, but the same equation can be obtained from a 2D model with the previously mentioned simplification and averaging across the direction orthogonal to the dominating flow (*y* axis).

Although substantial AP tissue flow occurs between 19 and 36 hAPF (peaking between 24 and 28 hAPF), we only study it in the vicinity of its maximum around 25 hAPF (Extended Data Fig. 1a). Since the tissue has already been flowing for several hours before our time of study, we assume that the transient regime due to tissue elasticity has vanished and that it can be considered as a purely viscous fluid. In other words, considering the tissue as a Maxwell material, for which the stress relaxes over the viscoelastic relaxation time (*τ* = *η/E, E* being the tissue elastic modulus), we study the tissue flow in the limit of long timescales, namely at times much larger than this relaxation time^10,11^. In this “long-time” limit, the tissue can therefore be considered as a viscous liquid and its elastic properties can be neglected. In the literature, one can find a wide range of estimates for *τ* in living tissues, from a couple of seconds in the *Drosophila* notum^15^, to minutes in embryonic tissues^12^, to hours in cell aggregates^13,14^. Importantly, although considered a as purely viscous fluid, the tissue remains nevertheless active since it is made of living cells that can exert forces on their substrate to migrate toward the neck.

In what follows the tissue characteristic length *L* now corresponds to the notum length along the AP axis, namely the distance between the rear of the notum and the notum edge, right before the neck furrow. Note that, accordingly to our previous assumption, *L* hardly changes over time as the notum rear end and the neck furrow barely move along the AP axis during the studied period, even though the latter deepens substantially along *z*.

### 2.2 Dissipative terms

When an elementary piece of tissue (of same area *dS, regardless of the animal size*, or in 1D, of length *dX*) is moving, it experiences a viscosity as it deforms and when parts of it are sliding past to one another. This results in a viscous stress (homogeneous to a force per unit area in 3D, but to a force in 1D):

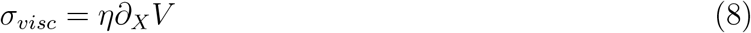

Note that in 1D, the shear and bulk viscous stresses are entangled, so the viscosity coefficient *η* is the sum of both coefficients.

Then, as the tissue moves, it also experiences friction from the apical extracellular matrix (aECM) at the top, onto which it actively crawls, and possibly from its cells attachment to the basal extracellular matrix (bECM) at the bottom. We write this term in the most simple possible way, as a force per unit length (since we writing it in 1D) proportional and opposed to velocity:

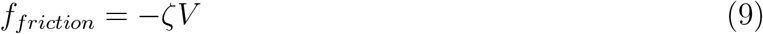

where *ζ* is the friction coefficient combining top and bottom frictions from the apical and basal ECMs.

### 2.3 Active term

Unlike passive materials that need external forces to deform them or make them flow, a developing tissue is an active material that can propel itself^1,16–18^. In our case, this activity enables the cells to crawl on the aECM, thereby resulting in the migration of the tissue toward the neck^2^. This migration, since it is not homogeneous in space, results in tissue deformation.

We assume that each cell exerts a maximum force −*ϕ* on the aECM thanks to the protrusions attached to it, and enabling them to crawl (Fig. 2c-f and Extended Data Fig. 2a-f). Considering the impact that impaired Dpy or polarity distributions have on the tissue flow, we assume that this ability to crawl is modulated by the combination of Dpy and polarity into an “activity” function *A*, for now unknown but bounded between −1 and 1, driving a migration to towards the neck when positive.

Since experimental observations indicate that the variations of Dpy and cell polarity along the AP axis scale with the animal size but keep the same amplitude, regardless of the animal size (Fig. 3a,c), we assume that *A* directly depends on *X/L*, thereby making its variations along the AP axis perfectly match the tissue size (Fig. 3f).

We therefore write the active force (per unit length of material) due to cell pulling on the aECM as follows:

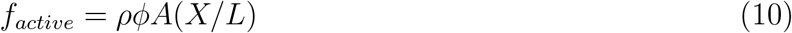

where *ρ* is the cell density.

The cell maximal active force *ϕ*, namely the ability of a cell to crawl depends on the amount of protrusions it is able to attach to the aECM. This amount of protrusions scales with cell apical size as experimental evidence show that protrusion are projected from a substantial part of the cell apex, and not just its periphery (Fig. 2c,e,j and Extended Data Fig. 2a,c,e,f). We therefore write:

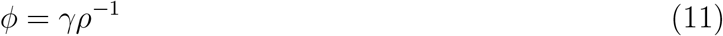

where *γ* is a force per unit length. The local cell density therefore vanishes and we obtain:

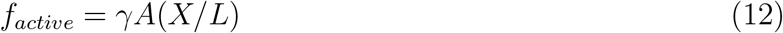

The local active force is then directly proportional to the activity function and therefore has its pattern scaling perfectly with tissue size (but not its amplitude, independent of tissue size).

### 2.4 Equation of motion

#### 2.4.1 Force balance

Since the Reynolds number is always much smaller than 1 in biological systems at this scale, we are in the overdamped regime and the inertial terms can be neglected when writing the force balance on an elementary piece of material, which leads to:

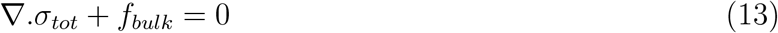

where *σ*_*tot*_ is the total stress exerted on a piece of material and *f*_*bulk*_ the total force acting on the bulk of the material, namely at each point of the material. Here, in 1D, we have: *σ*_*tot*_ = *σ*_*visc*_ (as we neglect the pressure term (see 2.6.4)) and *f*_*bulk*_ = *f*_*friction*_ + *f*_*active*_, which leads to the following equation of motion:

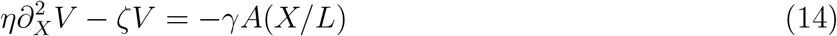

where we used the notation ∂_*X*_ = ∂*/*∂*X*. Dividing all terms by *η* leads to the appearance of the characteristic hydrodynamic length:

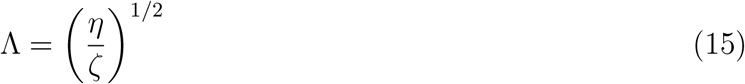

that determines the length scale over which the tissue responds to a mechanical perturbation. Eq. 14 rewrites:

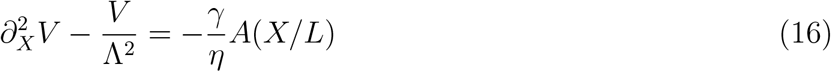

Thus, this is the equation that determines the velocity *V* of a given piece of tissue located at distance *X* from the rear of the notum, *regardless of the animal size*. This element of tissue moves thanks to the acto-myosin powered active force that gets dissipated by viscous and friction forces^16–19^.

Note that active parts of the tissue will influence passive parts of the tissue over distances comparable to Λ. Thus, some parts of the tissue that may locally experience little active force (for instance if *A*(*X*_*o*_*/L*) is small at some location *X*_*o*_) may still be moving at a substantial speed because of tissue viscosity, as it gets dragged by the other active parts of the tissue.

At this point, and as mentionned at the beginning, a same *X* value corresponds to different relative positions in tissues of different sizes. In the following, we will extensively use the dimensionless coordinate *x* = *X/L*, and a given *x* will therefore correspond to the same *relative* location in all tissues, regardless of their sizes.

#### 2.4.2 Dimensionless Equation

It is convenient and meaningful to rewrite the latter equation with dimensionless quantities, and to make the rescaled velocity *U* appear. We now rewrite it with variable *x* = *X/L*, which leads to:

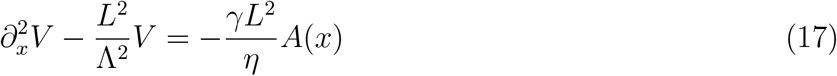

and the tissue length *L* now appears explicitly in the equation. Dividing the latter equation by *L* yields:

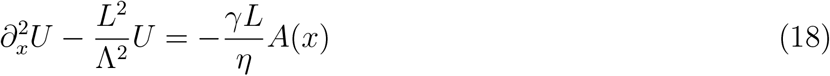

We now have an equation on *U*, but importantly, *L* still explicitly appears in this equation, making the observed linear scaling of velocity with *L* non trivial. Since the physical behavior of this active material is set by the characteristic length Λ, it will be relevant to express *L* as a multiple of Λ and therefore introduce the dimensionless tissue length *l* = *L/*Λ. By making it appear in the right hand side of the equation, we also make a characteristic rescaled velocity *U*_*o*_ appear (since *l* and *A*(*x*) are dimensionless):

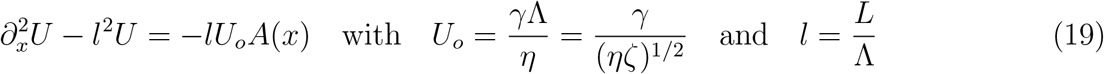

Instead of using *U*_*o*_ to renormalize *U*, which would lead to a quantity whose maximum value could be anything at this point, we will rather use a reference value of the rescaled velocity corresponding to the maximum value reached in the archetypal tissue of *L* = 656 *µ*m, namely:

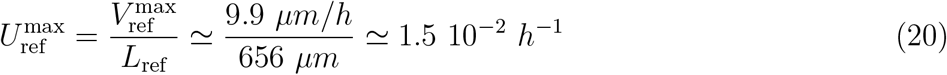

We therefore use 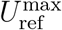 to normalize *U* and define the dimensionless rescaled velocity *u*:

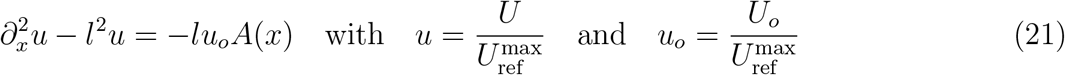

which is equation we must solve. Similarly, it can be convenient to define the dimensionless velocity *v*, that we define as follows:

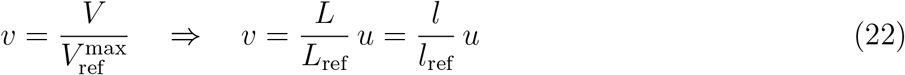

Note that the dimensionless velocity *v* is not simply *l u* but that it also involves *l*_ref_. Lastly, it is important to note that, given the relationship between *U* and the quantity 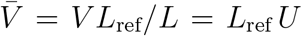 that we use throughout the main text (eq. 2), one has:

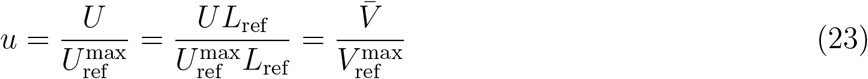

the latter relationship shows that this is strictly equivalent to plot *u* or 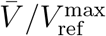.

### 2.5 Boundary Conditions & Limit Cases

#### 2.5.1 Boundary conditions

At the rear of the notum (*x* = 0), the tissue is fixed and does not flow, so we set:

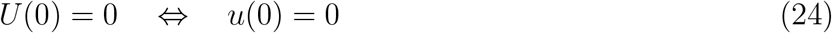

We apply the other boundary condition (BC) at the other end of the notum, near the neck, in the vicinity of *x* = 1. For simplicity, we assume that the neck constriction rates also linearly scale with the animal sizes. The rescaled velocities of the different tissue size bins then converge to a common value *U*_neck_ somewhere beyond the notum edge (*x* = 1) and the neck furrow that invaginates (*x >* 1), and we take it at *x*_*neck*_ ≃ 1.1. Note that the choice of alternative BC at the neck has little impact on the main results that we derive in the following, as will be discussed in section 2.6.3. At *x*_*neck*_, the tissue flow is therefore assumed to linearly scale with tissue size:

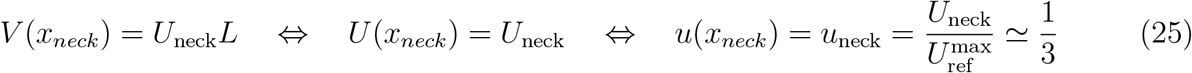

where *U*_neck_, and therefore *u*_neck_, are independent of *l* in good approximation. *u*_neck_ value can be estimated from the extrapolation of experimental velocity profile of the archetypal tissue along the AP axis up to the beginning of the neck furrow, leading to value 1/3 for control animals.

#### 2.5.2 No global scaling of tissue flow

Note that eq. 21 does not admit solution corresponding to global scaling of tissue flow at all scales. Indeed, injecting *u*(*x, l*) = *α*(*x*) into 21, one gets:

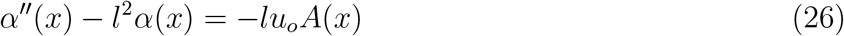

which must hold for any *x* ∈ [0, 1] and l ≥ 0. Taking the derivative of Eq. 26 with respect to *l* twice, yields:

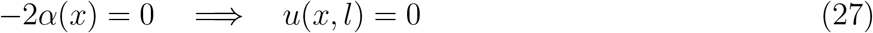

and there is no non-zero solutions of the form *u*(*x, l*) = *α*(*x*) that would enable scaling of velocity at all sizes. One therefore cannot achieve scaling at all scales in this framework, but local or approximate scaling may still be possible.

#### 2.5.3 Limit case 1: viscosity dominates: *l* ≪ 1 ⇔ *L* ≪ Λ

In this limit case the tissue is much smaller than the characteristic hydrodynamic length Λ. Eq. 21 therefore reduces to:

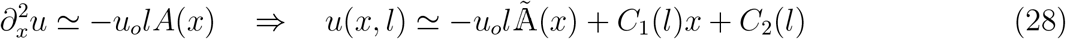

with

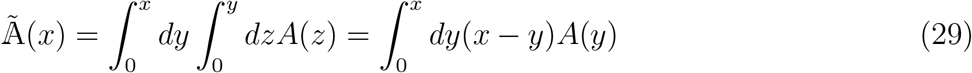

after integrating by parts. Using both boundary conditions leads to *C*_2_(*l*) = 0 and 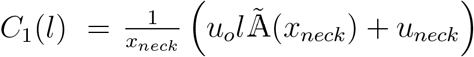, which in turn leads to:

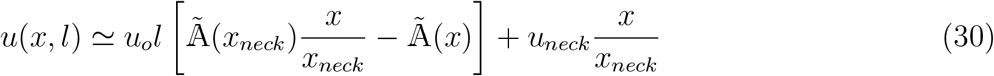

Note that, in this limit, the rescaled velocity *u* expression also displays a coupling between functions of *x* and *l*, precluding global scaling at all scales. In addition, *u* this time displays a linear dependence in *l* and therefore does not have an extremum in *l* either, precluding any kind of scaling.

Accordingly, in this limit, the velocity *V* (*x*) therefore reads:

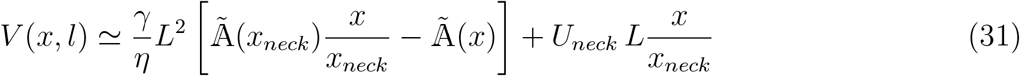

and *V* depends on *L*^2^ and animals are therefore expected to display a quadratic scaling of velocity amplitude with their length *L* (except near the neck given the BC we imposed at *x*_*neck*_), but not the linear scaling of velocity amplitude observed experimentally.

#### 2.5.4 Limit case 2: friction dominates: *l* ≫ 1 ⇔ *L* ≫ Λ

In this limit case the tissue is much larger than the characteristic hydrodynamic length Λ. Eq. 21 therefore reduces to:

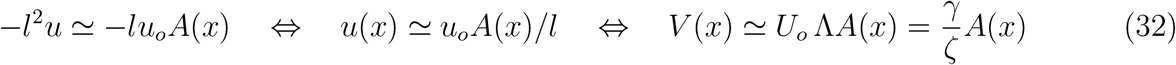

and the velocity *V* perfectly matches the active force profile, except close to the boundaries where boundary conditions must be met. The maximal velocity at zero viscosity therefore reads:

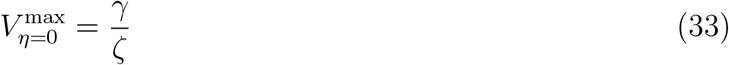

Note that, in this other limit, the rescaled velocity *u* expression displays a coupling between functions of *x* and *l* and therefore does not enable global scaling at all scales. In addition, *u* displays a dependence in 1*/l*, therefore decreasing with *l* and not having an extremum in *l* either, thereby precluding any kind of scaling.

Accordingly, in this limit, the velocity *V* (*x*) becomes independent of *L* and all animals are therefore expected to display similar velocities regardless of their size: there would be no scaling of velocity amplitude in this limit.

### 2.6 Remarks

#### 2.6.1 Scaling of velocity

From section 2.5.2 we know there will be no solution enabling global scaling at all scales. However, from those two limit cases where the rescaled velocity *u* features a regime where it increases with *l* (*l* ≪ 1 case), and a regime where it decreases with *l* (*l* ≫ 1 case) one can deduce that the general solution will at least admit a maximum in *l*, at a length we will call *l*^∗^, that may depend on *x*. Near this maximum, the general solution of *u* will therefore feature a local or approximate scaling of flow velocity where *u* is locally independant of *l*, or equivalently, *v* proportional to *l*, thereby describing the experimental observations.

#### 2.6.2 Physical parameters *η, ζ, γ*

It is important to note that the physical quantities *η, ζ* and *γ* represent the local intrinsic mechanical properties of the tissue, regardless of animal size. Thus, when considering a given elementary piece of tissue made up by a few cells to write the equations, we assume that these physical parameters remain the same regardless of the size of the animal. In other words, and similarly to what is done in fluid mechanics, we assume that the size of the system does not affect its local mechanical properties.

In addition, and for simplicity, we also assume that these parameters are homogeneous across the AP axis, namely that the physical properties of the tissue are the same across the AP Axis. Interestingly, assuming the parameters all depend on their position along the AP axis *x*, namely that they write 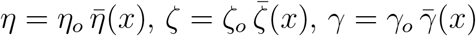 (“bar” quantities being dimensionless), will not change the conclusions obtained in the two limit cases. Indeed, eq. 21 then reads:

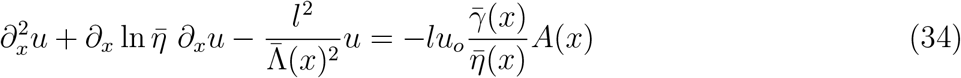

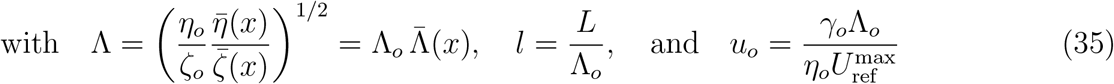

In each limit case 1 & 2, *u* will therefore still display a *l* and a 1*/l* dependence, respectively, thereby admitting a maximum in between, that corresponds to a linear scaling of velocity with *l*. However, regarding the variations with *x* in general, *u*(*x*) profile will obviously change in this case, as well as the *l*^∗^ dependence on *x*.

#### 2.6.3 Alternative Boundary Conditions (BCs)

To assess the robustness of our results to alternative BCs near the neck at *x* = 1, we tested different ones (not shown). Each time, we reploted the predicted velocity profiles in control tissue, laser cauterisation experiments and in the mutant conditions and compared it to the ones obtained with previous BC developped here.

The first alternative BC we considered near the neck was to directly impose the rescaled velocities measured experimentally at the neck boundary located at *x* = 1, namely *u*(1, *l*) = *u*_*exp*_(1, *l*).

The other alternative BC we considered was to, instead of setting *u*(1, *l*), alternatively set ∂_*x*_*u*(1, *l*), namely the slope of *u*(*x, l*) at *x* = 1. Physically, ∂_*x*_*u*(1, *l*) represents the (dimensionless) viscous stress near the neck at *x* = 1.

We find very similar results regardless of the BCs used. This illustrates the robustness of our results to the BC near the neck. In each case, for *l*_ref_ ≃ 3 and *u*_*o*_ ≃ 9 we find a nearly perfect scaling for 0 *< x <* 0.7 and some discrepancy for *x >* 0.7 following the trend observed experimentally. With these values, we also reproduce the tissue velocity profile observed in cauterization experiments and in mutant conditions.

#### 2.6.4 2D model assumptions

As mentioned before, this model can be written in 2D, which is more realistic, and lead to the same equation over *V*_*X*_ (eq. 14) by averaging along the ML axis (*Y* axis) and by making the following assumptions:

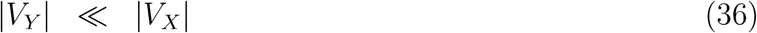

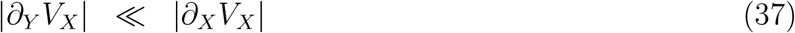

namely that the tissue flow along the transverse direction (*V*_*Y*_) is negligible with respect to the flow along the AP axis (*V*_*X*_), and that *V*_*X*_ variations in the transverse direction (*Y* axis) are negligible with respect to its variations along the AP axis (*X* axis), two assumptions that clearly holds when assessing the tissue flow field in the medial region (Fig. 1b,1f).

Lastly, more generally, a term of pressure appears in the isotropic part of the total tissue stress:

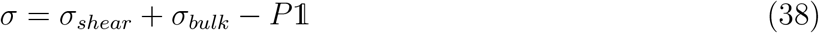

which, after averaging along the *y* axis and use of the above assumptions, leads to:

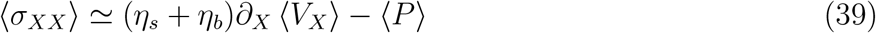

where *η*_*s*_ is the tissue shear viscosity and *η*_*b*_ the tissue bulk viscosity, that simply add up here to define *η*, as mentioned earlier (see 2.2).

In regular passive incompressible liquids, pressure gradients can drive their flows. Here, the cell active traction force is driving the flow, and friction and viscosity are mostly dissipating this force. Moreover, in 2D, cells are very compliant and their sometimes substantial deformation do not seem to limit the flow in any way. For these reasons we therefore neglect the pressure gradient term with respect to the viscous and friction terms, which then leads to the equation:

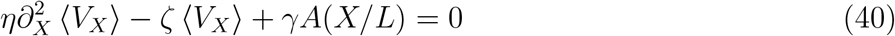

which corresponds to eq. 14, with ⟨*V*_*X*_⟩ renamed as *V*. Note that this derivation was achieved without the common assumption that the tissue is incompressible in 2D (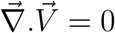 when incompressible), and that in 2D, each term has now the dimensions of force per unit area.

#### 2.6.5 Qualitative understanding of main equation

This equation 40 (or its 1D equivalent eq. 14) was derived implicitly considering a given elementary tissue area *dS*, regardless of the animal size, the same way we consider a given elementary volume of fluid *dV* when establishing the Navier-Stokes equation for Newtonian fluid, independently of the overall size of the system.

Nevertheless, things are a bit different here since the surface element will only contain a few cells, and that these cells are the ones generating the forces that drive the tissue flow. Since we know from the start that we have a range of animal sizes it is tempting to wonder how this change of size impact the dynamics of the tissue, not by considering a constant surface element or area *dS* regardless of the animal size, but precisely by *scaling* the surface element area with the tissue size. Obviously the results will not change, but it can be nevertheless informative to formulate it.

Let’s rewrite the equation 40 and multiply it by 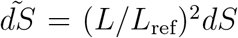, an elementary surface area that now scales with tissue size (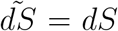 for reference tissue size *L* = *L*_ref_), to obtain three terms now having the dimension of a force:

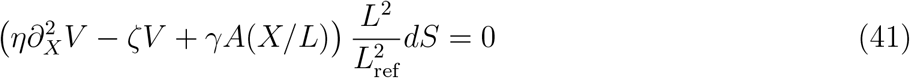

Now let’s distribute the (*L/L*_ref_)^2^ term in the parenthesis and use dimensionless *x* = *X/L* to “reset” all tissue coordinates in [0,1]:

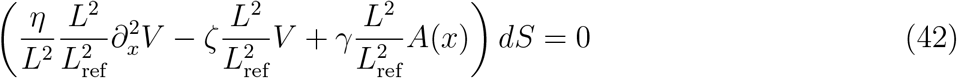

We now have an equation on the cell local velocity *V* (which is not dimensionless and is actually measured in *µm/h* in the lab referential) at relative location *x*. This equation is now written for surface area elements scaling with the tissue size.

Starting from the right, the active force term therefore scales with the tissue area since we now have (*L/L*_ref_)^2^ more cells exerting forces in the considered tissue element. Similarly, the friction term (2nd term) also increase with the tissue area (*L/L*_ref_)^2^. Thus, for friction dominated flows featuring only those two terms, the variation in tissue size does not change anything since both terms are changing in the exact same way and *V* is independent of *L* (see 2.5.4).

The viscosity term, on the other hand, behaves differently: it is also multiplied by (*L/L*_ref_)^2^ (since it was expressed per unit area like the other terms), but the second order space derivative makes a factor 1*/L*^2^ appear when resetting tissue spatial coordinates in [0,1] by using *x* instead of *X*. The space derivative therefore introduces a decreases in 1*/L*^2^ in bigger animals after rescaling their space coordinates.

The viscosity term dissipates energy when the velocity field display spatial variations, and it therefore favors homogeneous flows with cells moving together at similar speeds. Therefore, the same velocity variations in space measured in *relative* coordinates by 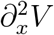 have a smaller influence in bigger animals because they correspond to lower actual velocity gradients. Indeed, this is well illustrated by the sketch of *A*(*x*) in Fig. 3f: the same *A*(*x*) profile in rescaled coordinates *x* (right panel) corresponds to weaker spatial variations leading to smaller gradients in bigger animals (left panel).

So the *L*^2^ increase due to higher surface area is compensated by the 1*/L*^2^ coming from the second order derivative, and as a result this viscosity term remains unchanged when tissue size varies. Thus, for viscosity dominated flows featuring only this term and the active force term, the velocity scales with *L*^2^ (see 2.5.3).

Finally, when both dissipative terms have comparable amplitudes, the characteristic hydrodynamic length Λ = (*η/ζ*)^1*/*2^ (eq. 15) becomes meaningful. This sets a typical length (in *µm*) over which a mechanical perturbation propagates in the tissue. Consequently, for a perfectly scaling pattern of activity *A*(*x*), as sketched in Fig. 3f (left panel), bigger animals, for which Λ is relatively shorter, will display velocity profiles closer to *A*(*x*) profile as compared to smaller animals. In this context, the velocity AP profiles of animals of various sizes cannot perfectly superimpose. However, within this crossover regime we know that we can find a length *L*^∗^ at which *V* displays a linear scaling with *L* (see 2.6.1).

### 2.7 General solution

We now derive the general solution of eq. 21 for any activity function *A*(*x*). The solution of the homogeneous equation writes:

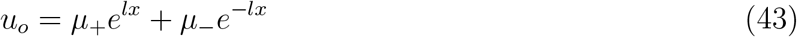

We look for a particular solution of the full equation 21 of the form:

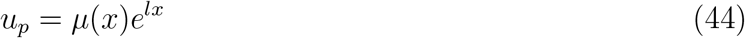

Injecting this expression into eq. 21, one finds that *µ* must satisfy:

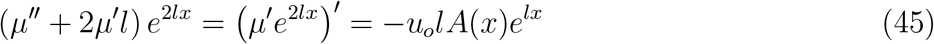

which must be integrated twice and yields:

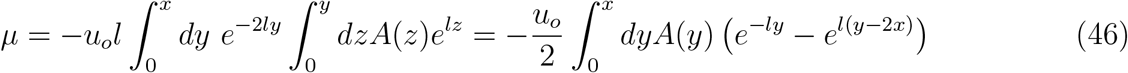

where the last equality was obtained by integrating by parts. This therefore leads to:

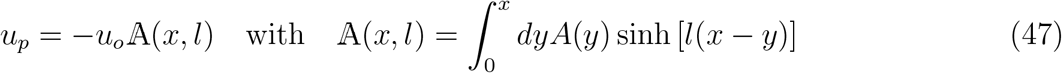

The general solution of eq. 21 therefore reads:

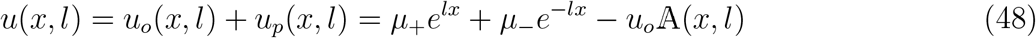

Now, satisfying both boundary conditions yields:

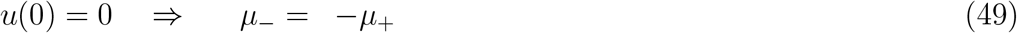

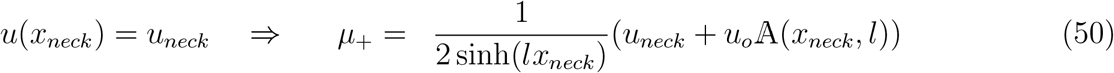

from which we get the general solution of eq. 21, respectively:

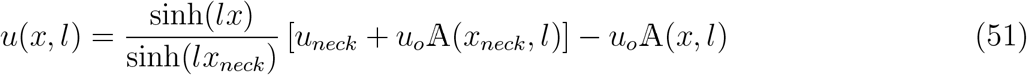

As expected, the general solution display a coupling between *x* and *l*, but as mentioned in section 2.6, it should reach a maximum enabling a scaling of velocity with tissue size where *u*(*l*) reaches its maximum.

In the particular case where the activity would be homogeneous across the AP axis, namely *A*(*x*) = *A*_*o*_, where *A*_*o*_ is a constant, A(*x, l*) then simplifies to:

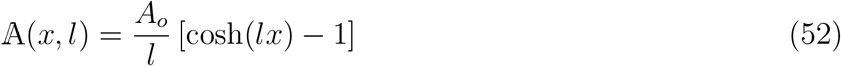

### 2.8 Activity profile *A*(*x*)

In order to determine and represent a rescaled velocity profile *u* satisfying eq. 21, we need to chose an activity profile *A*(*x*). Since Dumpy and cell polarity substantially impact the cell migration towards the neck, we write the activity *A*(*x*) as a simple combination of these two profiles along the AP axis:

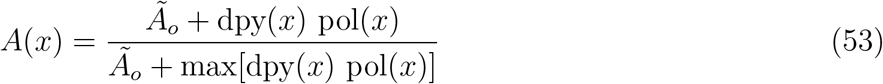

where dpy(x) and pol(x) are the Dumpy and cell polarity profiles, respectively, and *Ão* ∈ [0, 1] is a constant. *A*(*x*) thus defined is bounded in [0, 1]. We can rewrite it more conveniently as follows:

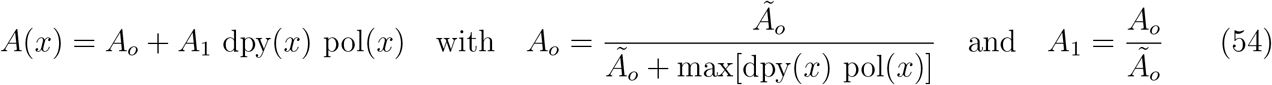

where *Ao* ∈ [0, 1] now represents an homogeneous activity across the AP axis accounting for the remaining migration observed in *dpy*^*RNAi*^ condition where dpy(*x*) ≃ 0 and therefore *A*(*x*) ≃ *A*_*o*_. In order to cover the full AP axis, we extrapolate the expression profiles to both bounds. The respective dpy(x) and pol(x) profiles, as well as the resulting combined profile are shown in fig. 4B and S4A-C.

### 2.9 Determination of velocity profile in control animals

#### 2.9.1 Adjustable parameters

There are therefore three adjustable parameters in our model:

- Λ, the characteristic hydrodynamic length, controls the spatial extent of the tissue response flow to applied forces. It is directly linked to the dimensionless tissue length *l* = *L/*Λ, and it will be set so that *l*_ref_ ≃ *l*^∗^, namely close to *u*(*l*) maximum. This will set the ratio *ζ/η* = 1*/*Λ^2^ ≃ (*l*^∗^*/L*_ref_)^2^.
- *u*_*o*_ that determines the amplitude of the flow and that will be set so that *u* reaches 1 at its peak for the “reference” tissue (archetype). This will therefore set the ratio 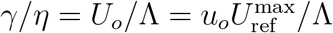.
- *A*_*o*_, the homogeneous activity base level across the AP axis, which influences the velocity profile along the AP axis. It determines the remaining flow in mutants where Dumpy levels and/or cell polarity are both substantially impacted since *A*(*x*) ≃ *A*_*o*_ then.

Note that the additional parameter *u*_neck_ corresponds to a boundary conditions and is chosen to extrapolate the experimental tissue flows at *x* = *x*_*neck*_. We set it to 1*/*3 for control animals and the mutant conditions (see section 2.5.1), except for *dpy*^*RNAi*^ and *dsh*^1^*sr*^1^ animals, which have substantially weaker flows, where it is set to 1/10.

#### 2.9.2 Predicted velocity profile *u*(*x, l*)

We now first explore the general solution by representing *u*(*x, l*) along the AP axis for different values of *l* and along the *l* axis, for different values of *x*.

Since we don’t know the values of the parameters relevant to our study yet, we systematically adjusts *u*_*o*_ so that *u* reaches 1 at its maximum, which leaves two free parameters Λ and *A*_*o*_. To evaluate the role of Λ (via the dimensionless tissue length *l* = *L/*Λ) we first need to assign a value to the baseline activity *A*_*o*_. Since the mutant conditions with the most impaired flows (namely *dpy*^*RNAi*^ and *dsh*^1^*sr*^1^) still display a rescaled flow that is roughly around 0.2 on average along the AP axis, we set *Ã*_*o*_ = 0.2 (⇒ *A*_*o*_ = 0.18) and represent *u*(*x, l*) for values of *l* ranging from 0.1 to 20 (Extended Data Fig. 4d). Note that for *l* = 20, which gets towards the *L* ≫ Λ limit and therefore a flow profile getting closer to the activity profile, one can check that this *Ã*_*o*_ value indeed results in a *u* plateauing around 0.2 in the rear part of the animal (Extended Data Fig. 4d).

As expected from the limit cases analysis and section 2.6, *u*(*x, l*) reaches a maximum in *l* at *l*^∗^(*x*) for any position *x* along the AP axis, with *l*^∗^(*x*) mildly depending on *x* in a non monotonous way (Fig. 4c,d and Extended Data Fig. 4e,f). This corresponds to the situation described in section 1.4 where the scaling, which occurs in the vicinity of *u*(*l*) maximum, can only be approximate. Indeed, the range of tissue sizes cannot simultaneously be centered on *l*^∗^(*x* ≃ 0.2) ≃ 2.5 at the rear of the notum, and centered on *l*^∗^(*x* ≃ 0.9) ≃ 3.4 at the front of the notum because of this variation with *x*. Therefore only some parts of the notum will optimally scale with its size, while the other parts will depart from optimal scaling. For this approximate scaling to occur, the physiological range of tissue sizes therefore need to lie in the vicinity of this range of maxima, namely *l* ∈ [2.2, 3.4] (Extended Data Fig. 4e).

#### 2.9.3 Determination of control model parameter values

Since *l*^∗^ depends on *x* and one cannot achieve a perfect scaling of flow velocity along the whole AP axis, the question of optimally choosing *l*_ref_ (or equivalently Λ) arises. Shall we favor scaling of velocity rather at the rear or rather at the front on the AP axis near the neck? Which tolerance should we set?

To determine the optimal value to assign to *l*_ref_, we evaluate the quality of the scaling by quantifying the area between the *u*(*x*) curves of the smallest and biggest tissue size bins. Since the experimental *u* curves overlay quite strikingly on the interval *x* ∈ [0, 0.7] (Fig. 4f), thereby displaying a scaling of velocity with good accuracy, we therefore use this interval to estimate this area as a quantification of the scaling accuracy:

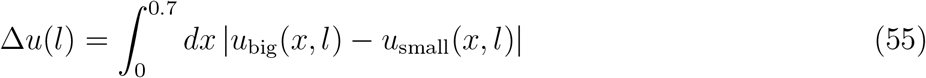

where *u*_big_ and *u*_small_ represent the rescaled velocity of the biggest and smallest tissue length bins, respectively, given a reference tissue length *l* (Extended Data Fig. 4g). The minimum of this function will therefore determine the best scaling we can obtain on this interval and will help determine *l*_ref_. We therefore represent Δ*u*(*l*) which reaches a minimum in the vicinity of *l* ≃ 3 (Extended Data Fig. 4h, blue curve). Since the scaling of experimental *u* is not perfect, we also calculate the experimental value of Δ*u*_*exp*_ ≃ 0.015, which sets the tolerance level (Extended Data Fig. 4h, purple straight line). Note that because the bin containing the smallest animals behaves substantially differently, we estimate Δ*u*_*exp*_ with the second smallest size bin rather than with the smallest one. Even with this precaution that lowers Δ*u*_*exp*_ value, Δ*u*(*l*) minimum is below the experimental value, meaning that the accuracy of the scaling reached theoretically is slightly better, which can be partially due to the absence of noise in the model. We therefore pick *l*_ref_ = 3 for simplicity, which is close to optimizing the superposition of the curves of the different tissue length bins in the interval *x* ∈ [0, 0.7] (Fig. 4g), like observed experimentally (Fig. 4f).

At this value, the curves superimpose almost perfectly in the *x* ∈ [0, 0.7] range, then, as expected since we know we cannot reach a perfect scaling on the whole AP axis, the curves separates for 0.7 ≲ *x* ≲ 1, with smaller animals displaying lower rescaled velocities *u* than bigger animals, as observed experimentally (Fig. 4f). Note that the lowest velocity curve that detaches from the other curves (purple) corresponds to the last set of viable animals whose morphogenetic processes, including migration to the neck, may be somewhat impaired. For values of *l*_ref_ outside of this range, which either corresponds to regimes where viscosity starts to dominate (*l*_ref_ < 3, before Δ*u*(*l*) minimum), or for regimes where friction starts to dominate (*l*_ref_ > 3, after Δ*u*(*l*) maximum), the rescaled velocity curves no longer superimpose (Extended Data Fig. 4i,j).

We therefore found a set of parameters that enables superposition of rescaled velocity curves *u*, thereby reproducing experimental observations with good approximation. This corresponds to a velocity *V* scaling linearly with tissue length *L*. We can therefore also estimate the characteristic length to be Λ = *L*_ref_*/l*_ref_ ≃ 656*/*3 ≃ 220 *µm*. Λ value therefore also sets the ratio *ζ/η* ≃ 2.1 10^−5^ *µm*^−2^.

With *l*_ref_ set, *u*_*o*_ is also set so that the reference tissue reaches a maximum rescaled velocity of 1 along the AP axis. We found *u*_*o*_ ≃ 8.6 and we can therefore also estimate the characteristic rescaled velocity 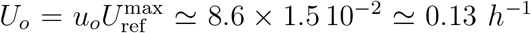. This also sets the ratio *γ/η* = *U*_*o*_*/*Λ ≃ 5.9 10^−4^ *µm*^−1^*h*^−1^.

By combining these two ratios, one can eliminate the tissue viscosity *η* to determine the ratio *γ/ζ* ≃ 28 *µm/h* which corresponds to the theoretical maximum velocity reached when the tissue viscosity *η* vanishes, namely 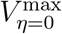 (eq. 32). This therefore leads to the corresponding dimensionless velocity 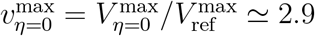.

We then compare the control average rescaled velocity profile with the one from the model including the standard deviation coming from the activity *A*(*x*) (Fig. 4b). The superimposition of the two display good agreement (Fig. 4h).

We can also check that the data points of the experimental maximum velocity versus tissue size overlay with the curve of predicted velocity and its deviation. As expected, the vast majority of data points lie within the expected range, with only the smallest animals slightly departing from the prediction (Fig. 4i). Also note that in the vicinity of 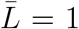 (namely *L* = *L*_ref_ or *l* = *l*_ref_ or 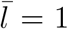), one has *u*(*x*^∗^, *l*) ≃ 1 (*x*^∗^ being the point on the AP axis where *u*(*x, l*_ref_) reaches 1, its maximum), and therefore (from Eq. 22):

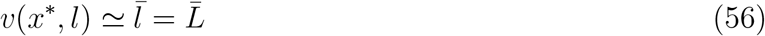

namely *v* becomes directly proportional to tissue length, as expected (dashed straight line in Fig. 4i).

### 2.10 Predicted *u* profiles in control tissue ablated conditions

All parameters have been set in the previous section. To further test the model, we now disturb the flow using laser cauterization to create obstacles near the neck or in the notum, therefore introducing an additional zero-flow boundary condition at in each case (Fig. 4j,k). We therefore apply these new boundary conditions in the equations and we solve them numerically. In both cases we find a decent agreement with the experimental curves, with the standard deviation (SD) of the predicted curves overlapping with the SD of the experimental curves, thereby confirming that the characteristic length Λ lie in the right range.

### 2.11 Predicted *u* profiles in mutant conditions

In order to further test the model, we then used genetic conditions that either disrupts Dumpy level (*dpy*^*RNAi*^), or polarity (*dsh*^1^), or both (*sr*^1^, *dsh*^1^*sr*^1^) (Fig. 2i,m,n).

For each condition, we renew the model prediction by replacing the control polarity and Dumpy profiles in *A*(*x*) (eq. 54) by the ones experimentally measured from each mutant conditions, thereby leading to multiple *A*(*x*) profiles matching each condition (Extended Data Fig. 4k). The model yields in each case a predicted velocity profile that we compare to the experimental curves (Fig. 4l-o). Given the simplicity of the model, we find an acceptable agreement between experimental and predicted curves, suggesting that the model, despite its simplicity, captures the essential elements of the process.

As already mentioned in section 2.9.1, *u*_*neck*_ was decreased from 0.33 to 0.1 for *dpy*^*RNAi*^ and *dsh*^1^*sr*^1^ conditions to better match the anterior limit of their experimental flows.

### 2.12 Predicted *u* dependence on physical parameters *η, ζ, γ*

Now that we have validated the model with different kinds of perturbations, we can use it to anticipate how, at a given size (the archetype reference length), the rescaled velocity will change with tissue viscosity *η*, external friction *ζ*, and cell active force *γ*. We plot the results in Extended Data Fig. 4l-n.

As expected, we find that decreasing viscosity or friction results in increasing the rescaled velocity (Extended Data Fig. 4l). However, each parameter has an opposite impact on the characteristic length Λ (eq. 15). Decreasing viscosity *η* will decrease Λ, leading to a *u* profile displaying a shape closer to *A*(*x*) (Extended Data Fig. 4l). In contrast, decreasing friction *ζ* will increase Λ, leading to a *u* profile displaying a shape further from *A*(*x*) (Extended Data Fig. 4m). Regarding the cell traction force *γ*, as expected, increasing it leads to increased migration velocity in the model (Extended Data Fig. 4n).

We can therefore also plot the predicted evolution of velocity *v* = (*l/l*_ref_) *u* as a function of *l* for changed viscosity *η* (Fig. 4s) or changed friction *ζ* (Extended Data Fig. 4o). Thus, when considering the physiological size range (roughly 0.5 *L*_ref_ − 1.1 *L*_ref_), the model predicts a decrease of the *V* (*L*) apparent slope (as we are then departing from the linear regime) if we manage to decrease the tissue viscosity *η* experimentally (Fig. 4s). Inversely, the model predicts an increase of the *V* (*L*) apparent slope in the physiological size range if we manage to decrease friction on ECMs *ζ* experimentally (Extended Data Fig. 4o). Note that in both cases the velocity increases as tissue flow is facilitated, but the *V* (*L*) trend evolves in opposite ways in each case.

### 2.13 Alternative friction: dependance on Dpy(x)

Since friction depends on attachment to the aECM and bECM, we investigated the possibility that the friction coefficient *ζ* may depend on Dpy level, which varies along the AP axis (Extended Data Fig. 4a). We therefore explored this possibility and defined 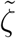 as follows:

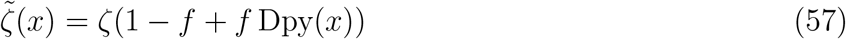

where *f* ∈ [0, 1] is a dimensionless parameter such that 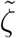 varies between *ζ* for *f* = 0, which correspond to the previously studied case, and *ζ* Dpy(*x*) for *f* = 1, this latter case corresponding to a friction directly proportional to Dpy profile. We therefore explored whether this added complexity improved the model by varying parameter *f*.

Taking *f >* 0 actually lead to decreased accuracy of the model for a simple qualitative reason: Dpy deposition being substantially decreased or suppressed in *dpy*^*RNAi*^, *sr*^1^, and *dsh*^1^*sr*^1^ mutant conditions, the friction 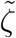 is then decreased in those mutant conditions for *f >* 0, which leads to increased velocity (Extended Data Fig. 4m), therefore leading to overestimation of *u* and deteriorated fit to the data as compared to Fig. 4l-o. We therefore kept the simple hypothesis of homogeneous friction along the AP axis, which corresponds to *f* = 0.

### 2.14 Conclusion

We developed a simple model of an active tissue able to migrate on its apical substrate thanks to an active force directly depending on tissue polarity and Dumpy level. The only two dissipative forces opposing this migration are tissue viscosity and friction on external ECMs.

Despite its simplicity, this model helps better understand the role of mechanical properties in the scaling of tissue flow. It indeed captures the observed linear scaling of velocity with animal length, which, since the physiological size range of animals is set, occurs at a specific value of the characteristic length Λ that emerges from the dissipative features of the tissue (viscosity over friction). It is also in good agreement with control tissue flows disrupted with laser ablations, and with mutant conditions impacting tissue polarity and/or dumpy deposition, as well as mutant conditions modulating tissue viscosity such as *p120* and *zip*^*DN*^.

